# Spontaneous Hemodynamic Oscillations during Sleep Coordinate Directional CSF Flow in the Subarachnoid Space and Ventricles

**DOI:** 10.64898/2026.10.07.757422

**Authors:** Danlei Chen, Fuyixue Wang, Amelia Strom, Hannah Yun, Jonathan R. Polimeni, Zijing Dong, Laura D. Lewis

## Abstract

During sleep, cerebrospinal fluid (CSF) flow is enhanced and thought to facilitate clearance of metabolic waste from the brain. Human imaging has shown that sleep is associated with large waves of bulk CSF flow in the fourth ventricle; however, the CSF flow dynamics around brain tissue are unknown. Here, we investigated how CSF flow is organized across the subarachnoid space, a key site both for fluid exchange across cortical surface and for transport out of the cranium. We used an MRI technology at 7 Tesla that simultaneously measures cortical hemodynamics and slow subarachnoid CSF flow and applied this during sleep. We found that spontaneous infraslow hemodynamic oscillations were coupled to spatially structured, directional CSF flow throughout both ventricular and subarachnoid compartments. Hemodynamic oscillations and fourth-ventricle bulk flow were both associated with brain-wide CSF flow at different phases, indicating that they reflect a single coordinated CSF movement cycle. Compared to cardiac pulsations, a canonical driver of CSF movement, hemodynamic-driven CSF flow showed a distinct directional pattern and was smaller in instantaneous amplitude but produced substantially greater cumulative displacement and thus greater potential for transport within an oscillation period. Together, these findings show that hemodynamic oscillations during sleep are coupled to temporally coordinated and directional CSF flow throughout the ventricles and subarachnoid space, suggesting that sleep-dependent neuronal activity drives fluid movement across the human brain in a distinct spatial pattern.

## Introduction

Sleep is essential for overall health, and its disruption is implicated in a wide range of pathologies (Ju et al., 2014; Krause et al., 2017; Lewis, 2021; Parhizkar & Holtzman, 2025; Sabia et al., 2021). Decades of work have shown that non–rapid eye movement (NREM) sleep is accompanied by large-scale neural activity and coordinated hemodynamic fluctuations (Brown et al., 2012; Diekelmann & Born, 2010; Steriade et al., 1993), observed in fMRI as infraslow oscillations in cortical gray matter (Fultz et al., 2019; Horovitz et al., 2008; Özbay et al., 2019; Picchioni et al., 2011). Sleep is also closely linked to the dynamics of cerebrospinal fluid (CSF), the fluid that fills the spaces surrounding the brain and has long been known to carry metabolic waste away from the brain (Cserr et al., 1977; Iliff et al., 2012; Tarasoff-Conway et al., 2015). More recent rodent studies have found this clearance process is more active during sleep (Iliff et al., 2013; Jiang-Xie et al., 2024; Xie et al., 2013), driven by sleep-related physiological oscillations (Hauglund et al., 2025). Furthermore, disrupted CSF dynamics are implicated in neurodegenerative disease (Nedergaard & Goldman, 2020; Rasmussen et al., 2018), highlighting the need to understand CSF transport dynamics.

In humans, fMRI studies have shown that CSF bulk flow in the fourth ventricle during sleep is temporally coupled to infraslow fluctuations of activity in the cerebral cortex (Fultz et al., 2019; Gonzalez-Castillo et al., 2022; Picchioni et al., 2022), providing initial evidence that sleep-related neural and hemodynamic activity may drive large-scale CSF dynamics. What has not been investigated, however, is how infraslow hemodynamic activity during sleep modulates CSF flow through the larger CSF system, including both the ventricles and subarachnoid space. The subarachnoid space is the brain-wide, CSF-filled space between the arachnoid and pia mater and it lies immediately adjacent to the brain tissue where metabolic waste is produced. It is a critical interface for fluid exchange between brain tissue and CSF (Bedussi et al., 2017, 2018; Iliff et al., 2012; Xie et al., 2013) and for transport of waste out of the cranium (Ahn et al., 2019; Louveau et al., 2015; Ma et al., 2017; Ringstad & Eide, 2020). Sleep-related bulk flow in the fourth ventricle has been proposed to support these same processes (Fultz et al., 2019; Lewis, 2021), but whether and how the fourth ventricle bulk flow waves are linked to flow in the subarachnoid space is unknown. Investigating CSF flow in the subarachnoid space is therefore essential for understanding how sleep-related hemodynamic events shape CSF dynamics and transport across the brain. CSF is also continuously driven by cardiac pulsation, a well-established, potent driver of CSF motion (Battal et al., 2011; Wagshul et al., 2011), producing pulsatile flow synchronized with the cardiac cycle. Determining whether the cardiac- and hemodynamic-driven flow reflect distinct modes of CSF transport, or the same underlying transport process operating across different temporal and spatial scales, is therefore essential for understanding whether sleep-related hemodynamic oscillations provide a distinct route for CSF transport beyond that already supported by cardiac pulsation.

Subarachnoid CSF dynamics have not been measured during sleep primarily due to methodological limitations. Subarachnoid CSF flow is slow and occurs within narrow, anatomically complex spaces, making it difficult to image and quantify. Conventional phase-contrast MRI captures relatively fast CSF flow in structures such as the cerebral aqueduct or ventricles (aqueduct velocity ∼30–80 mm/s; Battal et al., 2011; Nitz et al., 1992) but is less sensitive to the slower flow in the subarachnoid space (<2 mm/s; Dong et al., 2023, 2025; Wang et al., 2024). Fourth-ventricle inflow measurements provide a useful index of bulk movement into the ventricular system but do not resolve CSF spaces outside the ventricles (Fultz et al., 2019; Williams et al., 2023). Both phase-contrast and inflow approaches have demonstrated substantial CSF flow along the head– foot axis, but these measurements have largely been confined to large CSF compartments with fast flow. The broader CSF system has been measured using diffusion-weighted and motion-sensitized MRI approaches (Harrison et al., 2018; Hirschler et al., 2025; Taoka et al., 2017; Wright et al., 2024); however, these methods characterize fluid mobility that combines both coherent and incoherent motion rather than quantifying directional velocity, which is necessary to estimate fluid transport. Thus, quantitative imaging that directly measures slow flow with high sensivitiy is needed to characterize directional CSF flow dynamics and related solute transport during sleep.

Here, we employed a recently developed slow-flow-sensitized phase-contrast imaging (SOPHI) acquisition (Dong et al., 2023; Wang et al., 2024; Wang, Polimeni, et al., 2026; Wang, Reese, et al., 2026) with echo-planar time-resolved imaging (EPTI; Dong et al., 2024; Wang et al., 2019) at ultra-high field (7 Tesla) to measure CSF dynamics during sleep throughout the ventricular and broader subarachnoid spaces. SOPHI utilizes slow-flow-sensitive velocity encoding with the multi-echo readout of EPTI, enabling direct measurment of directional slow CSF velocity with distortion- and blurring-free imaging of across multiple CSF compartments, including the brain-wide CSF flow pattern in the narrow subarachnoid space (Wang, Reese, et al., 2026). Importantly, because this multi-echo readout captures both phase and magnitude information within the same acquisition, it enables concurrent measures of CSF velocity and cortical BOLD fluctuations with enhanced sensitivity within the same runs (Wang, Polimeni, et al., 2026). This simultaneous acquisition is necessary to study spontaneous dynamics in sleep, where large hemodynamic events occur intermittently. Using SOPHI-EPTI, we imaged participants during a nighttime sleep opportunity beginning near their habitual bedtime to capture naturalistic sleep. Velocity encoding was set along the head–foot axis, the dominant direction of bulk CSF movement through the ventricular system and along the neuraxis (Battal et al., 2011; Dong et al., 2025; Fultz et al., 2019; Wang, Polimeni, et al., 2026; Williams et al., 2023). We found that spontaneous hemodynamic oscillations during sleep were coupled to spatially structured CSF flow with clear pattern of changes in direction (hereafter referred to as directional flow) throughout both ventricles and subarachnoid space. This flow was linked to fourth-ventricle bulk flow as part of a single coordinated cycle and, compared with cardiac pulsation, showed a distinct directional pattern with greater cumulative displacement.

## Results

### CSF flow dynamics modulated by spontaneous hemodynamic fluctuations

We first investigated whether spontaneous hemodynamic fluctuations during sleep are linked to CSF flow across both the ventricles and subarachnoid space. We used the SOPHI-EPTI sequence and imaged CSF flow the head–foot axis (i.e., velocity-encoded phase signal) simultaneously with BOLD signal (i.e., BOLD-sensitive magnitude signal) in 9 subjects across 17 scanning sessions (some subjects completed more than one session; see *Methods*). The participants rested in the scanner during a nighttime sleep opportunity starting near their habitual bedtime (mean start time ∼11:00 PM). To infer arousal state, participants were instructed to press a button alternately with their index and middle fingers approximately 1 s apart for as long as they remained awake. During sleep, large-amplitude and low-frequency (∼0.01–0.15 Hz) cortical BOLD oscillations appear, and the peaks of these oscillations are coupled to CSF inflow in the fourth ventricle suggested by prior studies (Fultz et al., 2019). We therefore examined CSF flow throughout the brain, relative to these global BOLD events. Because cardiac pulsation has a strong influence on CSF signals (Fig. S2; Movie S1), we first regressed out the cardiac-locked flow component using slice-timing information (see *Methods*). We identified spontaneous hemodynamic oscillations by detecting peaks of the global BOLD signal from cortical gray matter (Fig. 1B–C; *N* = 17) and examined whether CSF flow was locked to these hemodynamic peak events (Fig. 1C). We found that CSF flow velocity changed relative to the timing of the BOLD peak, both in ventricles and subarachnoid space; these events also induced velocity changes in cortical and subcortical gray matter compartments, likely due to tissue motion but with smaller amplitudes (Fig. 1D; Fig. S3A). Across subjects, whole-brain CSF flow showed significant changes, with the largest changes around the peak of the BOLD oscillation and a reversal of flow direction across the cycle (Fig. 1E; Movie S2), consistent with an oscillatory, spatially coordinated CSF movement. These time-locked changes were observed across both ventricular and subarachnoid spaces, indicating that sleep-related BOLD-locked CSF dynamics reflect a brain-wide, coordinated phenomenon rather than a localized one.

**Fig. 1.**
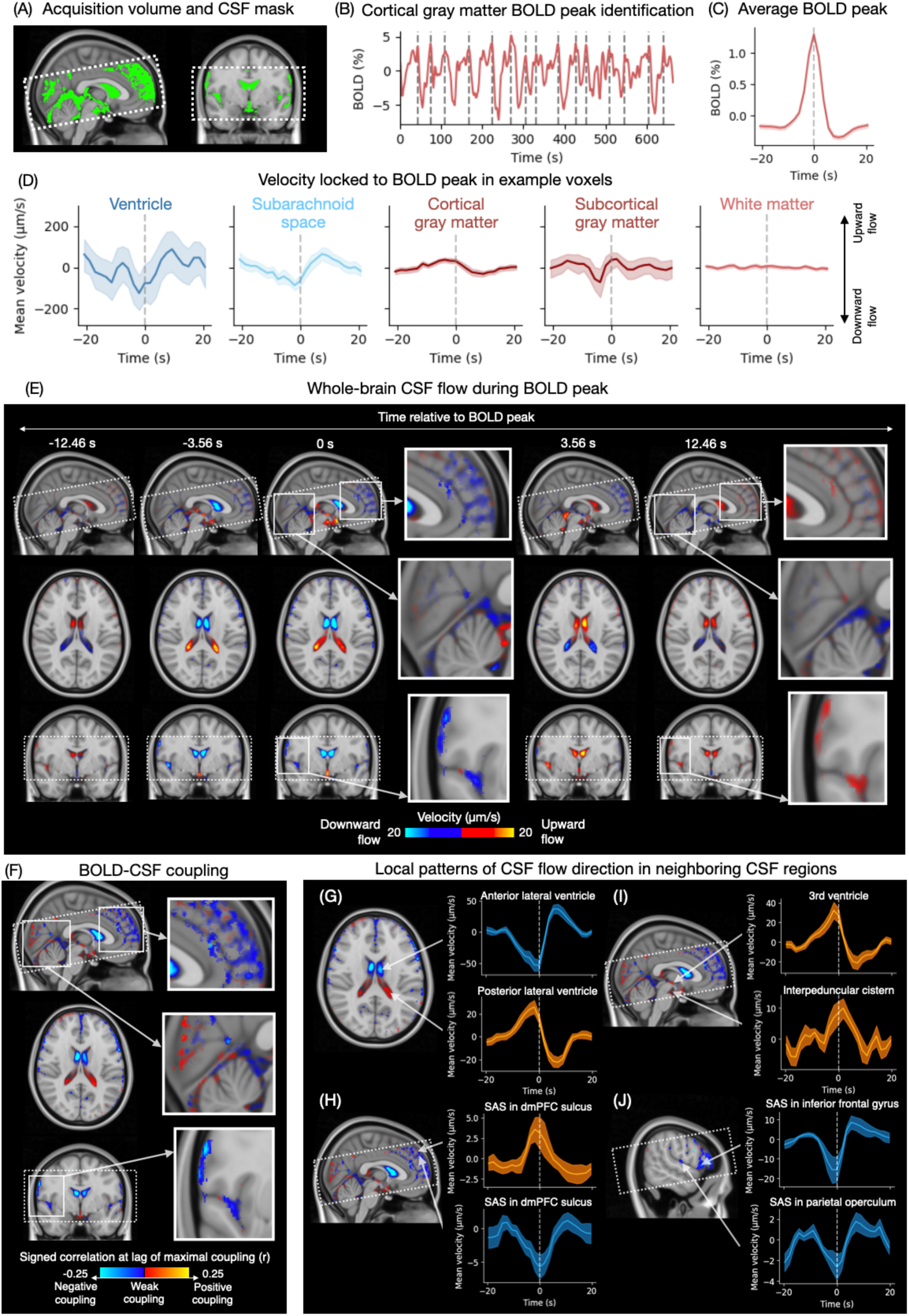
CSF flow velocity modulated by spontaneous hemodynamic oscillations. (A) Example acquisition volume and CSF mask. The acquisition volume, shown as dotted white box, was positioned to include the lateral and third ventricles and the surrounding mid-axial subarachnoid space. A group-level averaged CSF mask in green is shown for illustration purposes; main analyses were all conducted in subject native functional space. (B) Cortical gray matter BOLD peak identification. Spontaneous hemodynamic oscillations were identified as peaks in the run-level mean cortical gray matter time series. (C) Average BOLD peak time series locked to BOLD peaks, computed by averaging across all identified peaks from all scan sessions (*N* = 17 sessions; 9 unique participants). Error bar indicates 95% CI. (D) Example voxel time series showing CSF flow velocity and tissue motion time-locked to the BOLD peak, averaged across one representative run. Voxels in the ventricle and subarachnoid space both show CSF flow velocity changes around the BOLD peak, whereas voxels in tissue, including cortical and subcortical gray matter, show tissue motion around the BOLD peak; white matter shows relatively less tissue motion. (E) Whole-brain CSF flow across BOLD peaks shows spatially organized, oscillatory flow patterns time-locked to the BOLD peak. CSF flow dynamics time-locked to BOLD peaks were normalized and averaged across subjects. Yellow-to-red colors indicate upward flow, and blue voxels indicate downward flow. (F) BOLD–CSF coupling map. BOLD–CSF coupling is defined as the maximum absolute cross correlation between each voxel-wise time-locked CSF dynamic and the average run-level BOLD peak. Because the BOLD reference is a positive peak, positive BOLD–CSF coupling suggests a velocity change with a peak relative to its baseline, and negative coupling suggests velocity change with a trough. Larger values in either direction indicate stronger temporal coupling between BOLD peaks and CSF flow. (G–J) Local patterns of CSF flow direction in neighboring CSF regions. Brain maps show BOLD–CSF coupling. (G) BOLD-locked CSF velocity in anterior and posterior lateral ventricle voxels shows opposite-sign coupling, consistent with flow shifting back and forth within the ventricular compartment. (H) The opposite-sign pattern in adjacent subarachnoid space voxels in dorsomedial prefrontal cortex indicates local mixing within the subarachnoid space. (I) Third ventricle and interpeduncular cistern voxels both show positive coupling sign, indicating coherent flow along the basal outflow pathway. (J) Subarachnoid space voxels on the lateral surface near the inferior frontal gyrus and parietal operculum show negative coupling sign, indicating that spatially separated subarachnoid regions can also share coherent flow, possibly shaped by the local anatomy of connected CSF spaces. In panels G–J, orange traces indicate flow pattern with positive coupling sign; blue traces indicate flow patterns with negative coupling sign. Time series are shown as group averages, with error bars indicating 95% confidence intervals across scan sessions (*N* = 17). In all panels, dotted white box indicates acquisition volume placement.

To test whether the CSF flow pattern was reliably and significantly coupled to the BOLD signal, we computed the voxel-wise cross-correlation between the average flow dynamics in each CSF voxel and the average cortical BOLD peak within the same run. For each voxel, we extracted the maximum absolute cross-correlation across all lags; this measure was then compared to a null distribution obtained by phase-shuffling the CSF time series (preserving its power spectrum) and recomputing the cross-correlation with the original BOLD signal (see *Methods*), generating a voxel-wise significance estimate of BOLD–CSF temporal coupling. We observed strong BOLD–CSF coupling (voxel-wise permutation test, *p* < 0.05) for 30.57% of all CSF voxels (95% CI [23.98, 37.16]), including 32.22% (95% CI [24.60, 39.84]) subarachnoid space voxels and 25.53% (95% CI [21.79, 29.29]) ventricle voxels (Fig. S3B). Subarachnoid space voxels showed a greater proportion of significant coupling than ventricle voxels (paired *t*_(16)_ = −2.40, *p* < 0.05), suggesting that cortical hemodynamic changes may have a stronger influence on the subarachnoid space, which directly surrounds the cortex, than in the ventricular compartment. These results show that sleep-related hemodynamic oscillations are associated with CSF flow throughout the ventricles and the subarachnoid space, with the flow direction inverting around the hemodynamic peak, revealing a broad spatial scale of coordinated flow.

We next examined the spatial organization of flow direction in BOLD-driven CSF dynamics. The voxel-wise signed BOLD–CSF coupling map (i.e., maximum cross-correlation between each CSF voxel and the cortical BOLD peak; Fig. 1F) was used to capture the reliability of the dynamic coupling between BOLD and CSF flow. This measure is independent of each voxel’s absolute flow magnitude or baseline offset, allowing comparison across regions with very different physical flow scales. The sign of the BOLD–CSF coupling map reflects the direction of the relative velocity change at the lag of its maximum correlation: because the BOLD reference is a positive peak, positive coupling indicates a velocity rise then drop relative to its own baseline (i.e., peak; Fig. 1I), and negative coupling indicates a velocity drop then rise (i.e., trough; Fig. 1J). The spatial pattern of this coupling map closely resembled the velocity map at the BOLD peak (*r* = 0.72; Fig. 1F & Fig. 1E at *t* = 0 s), suggesting that the sign of the BOLD–CSF coupling, which reflects relative velocity change from each voxel’s own baseline rather than absolute flow direction, nonetheless closely tracked the actual direction of velocity at the BOLD peak (see Methods).

We observed spatially clustered patterns in the BOLD–CSF coupling map. For example, voxels in the lateral ventricles formed two clusters: an anterior cluster with strong negative coupling and a posterior cluster with strong positive coupling around the BOLD peak, such that their directional flow patterns were reversed relative to each other (Fig. 1G). This anterior–posterior reversal suggests a coordinated displacement within the ventricle, in which CSF is transiently moved from one end toward the other. A similar sign reversal was observed in neighboring subarachnoid regions located near the dorsomedial prefrontal cortex (Fig. 1H). This pattern of reversed oscillations demonstrates that CSF moves back and forth between adjacent connected compartments (i.e., local mixing) rather than in a single directional pattern. Conversely, some locations showed the same coupling sign, indicating they share the same direction of velocity change. For example, clusters in the third ventricle and interpeduncular cistern showed similar flow dynamics, both with strong positive coupling around the BOLD event (Fig. 1I), suggesting coordinated flow along a shared pathway. By contrast, the subarachnoid space in the inferior frontal gyrus and parietal operculum both showed strong negative coupling around the BOLD event (Fig. 1J). We also examined the timing of these dynamics, by calculating the temporal lag of the BOLD–CSF correlation. We also examined the timing of these dynamics, by calculating the temporal lag of the BOLD–CSF correlation. Although the coarse sampling rate (TR = 1.78 s) may not resolve finer temporal detail, CSF flow changes slightly led the BOLD event when averaged across significantly coupled subarachnoid voxels (lag = −0.77 s, 95% CI [−1.23, −0.32]), indicating that CSF flow changes were approximately concurrent at this temporal resolution.

Together, these results demonstrate that large hemodynamic oscillations during sleep coordinate brain-wide changes in CSF velocity. Furthermore, these CSF dynamics are spatially organized rather than uniformly distributed, showing structured, directional signatures across both ventricular and subarachnoid compartments. These directional patterns include reversed coupling between adjacent regions, reflecting local displacement or mixing within a shared CSF space, and consistent coupling along connected ventricular compartments with coherent, unidirectional flow.

### CSF flow dynamics modulated by fourth ventricle inflow

Flow-sensitized fMRI is widely used to detect CSF flow in the fourth ventricle (e.g., Fultz et al., 2019; Williams et al., 2023), which enables robust measurement of bulk CSF flow, and this measure shows large oscillations during sleep (Fultz et al., 2019; Picchioni et al., 2022). However, how this localized CSF inflow is related to flow in the subarachnoid space and the rest of the ventricles is not yet known. We therefore analyzed how distributed CSF velocities change relative to the oscillation in CSF inflow signals in the fourth ventricle. We first identified CSF inflow signal within the fourth ventricle from the bottom slices of the BOLD-sensitive magnitude images (Fig. 2A) and detected the peaks of this inflow signal (Fig. 2B), providing an index of bulk CSF inflow analogous to that used in previous fMRI experiments. The cortical gray matter BOLD signal was anticorrelated with the inflow peak (Fig. 2C & D; *N* = 16, as one of the sessions did not show reliable inflow effect), reflecting the established coupling between large-scale cortical hemodynamics and bulk ventricular CSF flow (e.g., Fultz et al., 2019).

**Fig. 2.**
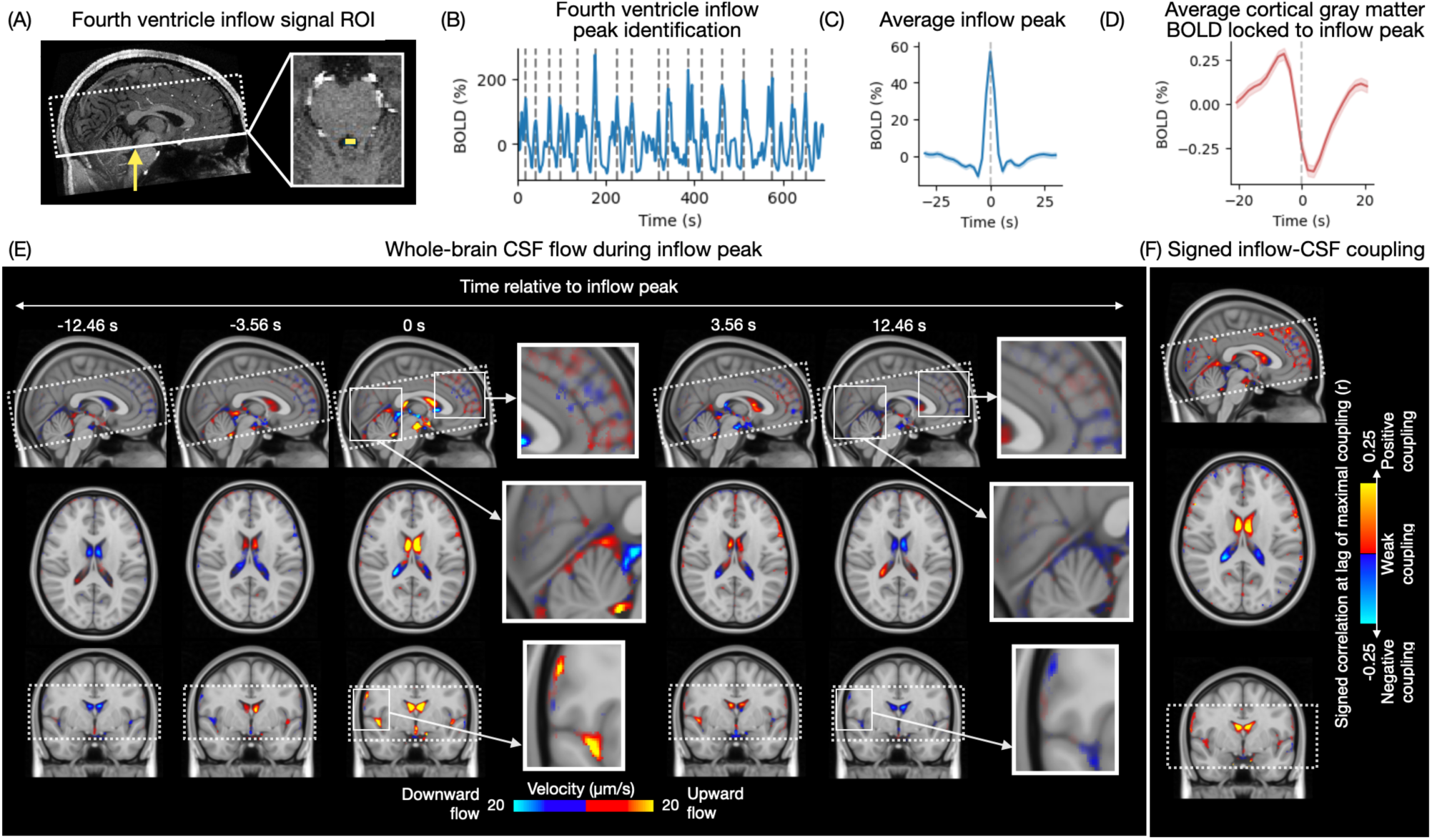
Large-scale CSF flow in the fourth ventricle is coupled to structured CSF flow throughout the ventricular and subarachnoid spaces. (A) Fourth ventricle inflow voxel identification. Inflow voxels were identified from high-intensity voxels in the bottom slices of the magnitude images. (B) Fourth ventricle inflow peak identification. Inflow peaks were identified as peaks in the run-level mean inflow time series. (C) Average inflow time series locked to inflow peaks, computed by averaging across all identified peaks from all subjects (*N* = 16 sessions). Shading indicates 95% CI. (D) Average cortical gray matter time series locked to inflow peaks. (E) Whole-brain CSF flow across inflow peaks showed spatially organized, oscillatory flow patterns time-locked to the inflow peak. CSF flow dynamics time-locked to inflow peaks were normalized and averaged across subjects. Yellow-to-red colors indicate upward flow, and cyan-to-blue colors indicate downward flow. (F) Signed inflow–CSF coupling. Because the reference inflow peak was always defined as a positive oscillation, in-phase CSF voxels with positive correlations showed positive oscillatory pattern, whereas anti-phase CSF voxels with negative correlations showed negative oscillatory pattern. Larger absolute correlation values indicate stronger temporal coupling between inflow peaks and CSF flow. SAS: subarachnoid space; GM: gray matter; WM: white matter. Dotted white box indicates acquisition volume placement.

Similar to the BOLD-locked flow patterns, inflow peaks were also accompanied by widespread, coordinated, and oscillatory velocity changes across ventricular and subarachnoid CSF (Fig. 2E; Movie S2; Fig. S4A & S4B). In particular, we observed strong upward flow in the fourth ventricle near the bottom slice at the peak of inflow event (middle panel of zoomed-in figure at *t* = 0 s in Fig. 2E), validating that the magnitude-based inflow signal reflects upward CSF movement into the fourth ventricle. In contrast to BOLD-locked results, a greater proportion of ventricle voxels showed significant coupling with inflow than the proportion in the subarachnoid space (permutation tests; all CSF spaces: Mean = 33.12%, 95% CI [26.54, 39.70]; Subarachnoid space: Mean = 29.08%, 95% CI [22.10, 36.07]; Ventricle: Mean = 45.02%, 95% CI [37.59, 52.45]; Subarachnoid space vs. ventricle: paired *t*_(15)_ = −7.81, *p* < 0.001; Fig. S4C), likely reflecting the anatomical proximity of the inflow signal to the ventricular system. To characterize the direction of inflow-locked velocity changes across the brain, we computed a signed maximum cross-correlation map to show coupling between inflow and CSF velocity (Fig. 2F). The sign of this inflow-locked map was largely inverted relative to the BOLD-locked map (Fig. 1F), consistent with inflow and BOLD reflecting the same underlying oscillation in different phases (Fultz et al., 2019; Williams et al., 2023). Together, these results demonstrate that fourth-ventricle inflow, previously used as a marker of localized bulk CSF motion, is coupled to coordinated flow across the ventricles and subarachnoid space.

### Relationship between BOLD-locked and inflow-locked subarachnoid CSF dynamics

Because cortical BOLD and fourth-ventricle inflow are themselves temporally coupled during sleep, the two reference events may index overlapping phases of the same CSF flow cycle. Further, whether the shared coupling is spatially uniform across subarachnoid space, or varies by region in ways that reveal distinct local contributions, is unknown. To address these questions, we compared BOLD- and inflow-locked flow patterns across subarachnoid regions. We first defined a subarachnoid space parcellation, by assigning each voxel the label of the majority tissue ROI directly surrounding it (Fig. 3A; 170 subarachnoid ROIs in total). We then averaged voxel-wise CSF flow within each subarachnoid ROI for the BOLD-locked and inflow-locked events (Fig. 3B; 109 ROIs (64.12%) and 126 ROIs (74.12%) that showed significant coupling to BOLD and inflow peaks, respectively), validating that ROI-level averaging captures robust coupling to both events. At the BOLD peak, subarachnoid CSF flow was on average downward (mean peak-to-trough velocity amplitude = 7.61 μm/s), consistent with the account that global BOLD peaks reflect increases in cortical blood volume through neurovascular coupling, which displace CSF out of the cranium given the fixed intracranial volume. In contrast, at the inflow peak, subarachnoid CSF flow was on average upward (mean velocity amplitude = 9.00 μm/s), consistent with CSF entering through the fourth ventricle and displacing subarachnoid CSF upward.

**Fig. 3.**
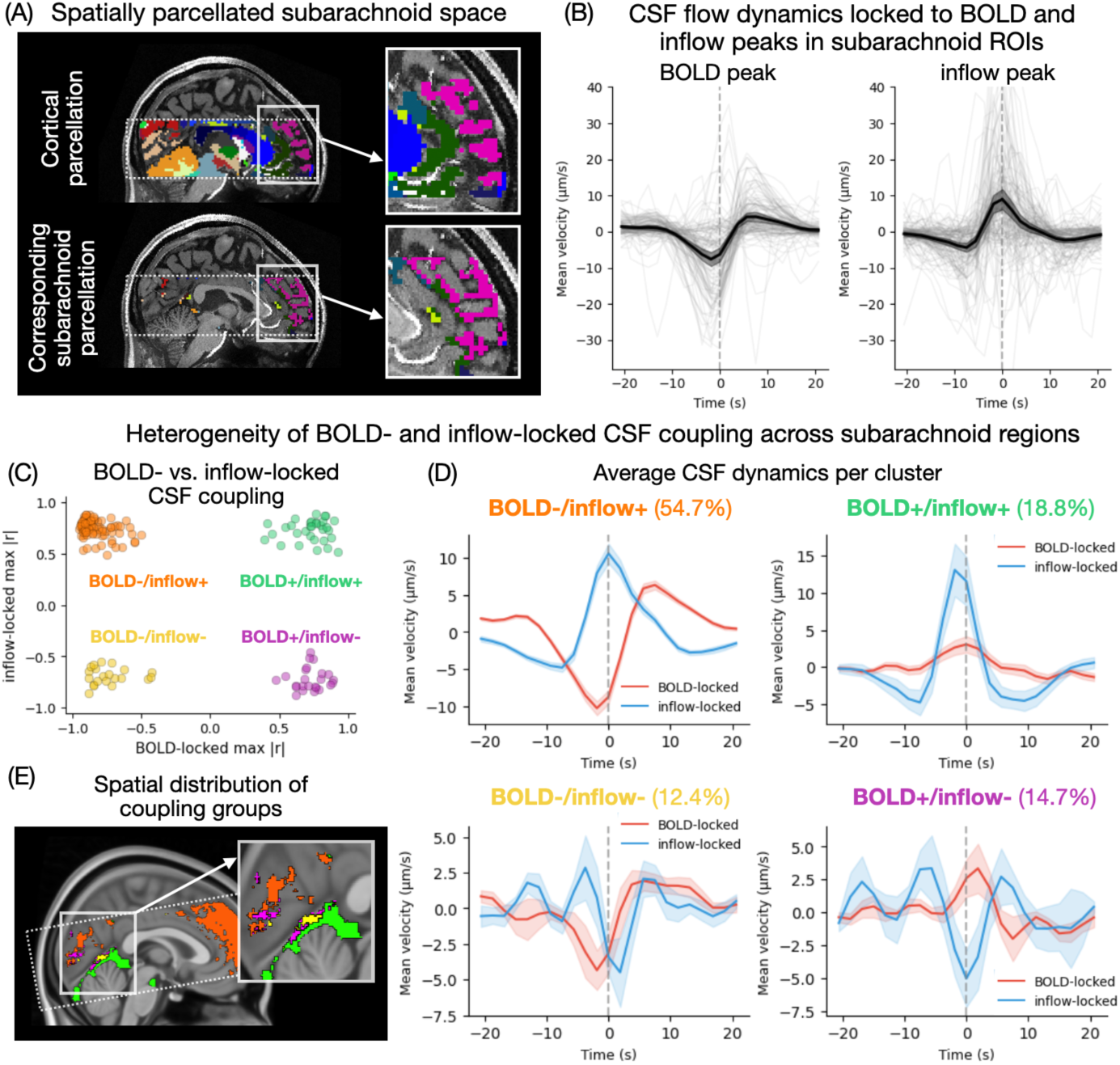
BOLD-locked and inflow-locked CSF dynamics reveal spatially heterogeneous CSF flow across the subarachnoid space. (A) Subarachnoid space parcellation, generated by assigning each CSF voxel the label of the majority surrounding tissue ROI in subject space. (B) Averaged CSF flow time-locked to cortical BOLD peaks and fourth ventricle inflow peaks in each subarachnoid ROI. Gray lines indicate the time series for each subarachnoid ROI that showed significant temporal coupling with BOLD and inflow peaks. Black lines with error bar indicate cross-subject average with 95% confidence interval. (C) Joint distribution of BOLD- and inflow-locked coupling across the subarachnoid ROIs. Each dot represents one ROI. (D) Group-averaged CSF velocity within each group, time-locked to BOLD peaks (red) and fourth-ventricle inflow peaks (blue). Groups are defined by the combination of BOLD-locked and inflow-locked coupling: BOLD−/inflow+ (negative BOLD-, positive inflow-locked coupling; 54.7% of all ROIs); BOLD+/inflow− (positive BOLD-, negative inflow-locked coupling; 14.1%); BOLD+/inflow+ (positive on both; 18.8%); BOLD−/inflow− (negative on both; 12.4%). Shaded bands indicate 95% confidence intervals across ROIs. (E) Spatial distribution of coupling groups across the subarachnoid space. Colors indicate groups defined in panel C. Dotted white box indicates acquisition volume placement.

We then tested whether BOLD- and inflow-locked CSF dynamics reflect the same underlying oscillation with a timing offset. For each subarachnoid ROI, we calculated cross-correlation between average BOLD- and inflow-locked flow dynamics. We found that the BOLD- and inflow-locked CSF flow were highly similar in oscillatory pattern (peak cross-correlation *r* = 0.81), with BOLD-locked CSF dynamics occurring 4 TRs (7.12 s) after inflow-locked dynamics. These spatially consistent yet temporally shifted whole-brain oscillatory patterns derived from these two independently identified signals indicate that subarachnoid CSF dynamics, ventricular inflow, and cortical hemodynamics are not separate phenomena but likely successive phases of a single brain-wide oscillatory process.

Although the ROI-level spatial anticorrelation indicates a global tendency for CSF flow to reverse between BOLD and inflow peaks, anatomical constraints, drainage pathways, and local vasculature could produce regional heterogeneity in how individual subarachnoid regions are organized. We therefore examined whether this BOLD- and inflow-locked pattern was consistent over regions. We grouped subarachnoid ROIs based on the sign of their BOLD- and inflow-locked coupling, yielding four groups with clear spatial organization (Fig. 3C). The most common group (BOLD−/Inflow+; 93 ROIs, 54.7%; Fig. 3D) showed negative BOLD- and positive inflow-locked coupling and was concentrated mostly along the dorsal cortical surface (Fig. 3E, orange; Fig. S5). These flow patterns showed similar absolute peak velocity (i.e., maximum absolute velocity across all lags) around BOLD and inflow peaks (BOLD: 10.64 μm/s; inflow: 11.55 μm/s; paired *t*_(92)_ = 0.83, *p* = 0.406; Fig. 3D), consistent with a symmetric reversal of flow between BOLD and inflow peaks and with the idea that global hemodynamic events more directly affect cortical-adjacent, wide-spread subarachnoid regions. A smaller set of ROIs (BOLD+/Inflow−; 24 ROIs, 14.1%; Fig. 3D) showed the opposite pattern, with negative inflow- and positive BOLD-locked coupling (absolute peak velocity: BOLD: 4.47 μm/s; inflow: 5.97 μm/s; paired *t*_(23)_ = 0.59, *p* = 0.558). These regions were spatially scattered without a clear anatomical clustering based on the current ROI definition (Fig. 3E, purple; Fig. S5). The positive-coupling group (BOLD+/Inflow+; 32 ROIs, 18.8%; Fig. 3D) at both BOLD and inflow peaks was concentrated in ventral basal cisterns near the brainstem and basal cortex (Fig. 3E, green; Fig. S5). For these ROIs, the absolute peak velocity was significantly greater for inflow-locked than for BOLD-locked (BOLD: 4.53 μm/s; inflow: 15.84 μm/s; paired *t*_(31)_ = 3.99, *p* < 0.001), indicating that flow in these basal regions is more strongly locked to bulk ventricular CSF inflow, consistent with their direct continuity with the fourth ventricle. Although the basal cisterns are major sites of CSF drainage, flow in these regions maintained the same direction around both BOLD and inflow peaks, suggesting that the sleep-related hemodynamic oscillation drives flow opposite to net drainage throughout the cycle, potentially facilitating CSF circulation and mixing. The negative-coupling group (BOLD−/Inflow−; 21 ROIs; 12.4%; Fig. 3D) was more scattered (Fig. 3E, yellow; Fig. S5; absolute peak velocity: BOLD: 4.83 μm/s; inflow: 6.52 μm/s; paired *t*_(20)_ = 0.76, *p* = 0.456), possibly reflecting a local driving force, such as proximity to large arteries or draining veins that impose a consistent local flow direction independent of the global BOLD–inflow cycle. These results suggest that while flow in most of the subarachnoid space is organized around the cortical hemodynamic cycle, a subset of regions follows distinct patterns, with basal drainage regions actively and consistently modulated by the cycle despite their net drainage flow and scattered regions possibly shaped by local vasculature, pointing to functional heterogeneity in how different parts of the subarachnoid space contribute to CSF circulation.

### Comparison between hemodynamic- and cardiac-driven CSF dynamics

Cardiac pulsation (∼1 Hz) is the canonical driver of CSF motion in human imaging, producing fast, rhythmic flow throughout the ventricles and subarachnoid space (Battal et al., 2011; Wagshul et al., 2011) with coherent brain-wide spatiotemporal patterns (Wang, Reese, et al., 2026). Whether the infraslow (∼0.05 Hz) BOLD-driven flow we describe above reflects a similar or fundamentally distinct mechanism is unknown. To address this, we directly compared CSF dynamics locked to the cardiac cycle and to spontaneous BOLD peaks.

We first visualized CSF flow dynamics locked to the cardiac cycle and to BOLD peaks for comparison. For voxels located in the same compartments, CSF dynamics differed across cardiac vs. BOLD oscillations in both flow amplitude and direction (Fig. 4A). For example, two voxels located in posterior and anterior portions of the lateral ventricle showed oscillatory cardiac-locked dynamics with opposite flow direction, whereas their BOLD peak-locked dynamics were more biphasic and also inverted in direction relative to each other (Fig. 4A). Two subarachnoid voxels located along the anterior medial wall showed similar cardiac-locked CSF profiles, but their BOLD peak-locked dynamics differed in flow direction (Fig. 4B). Likewise, voxels along the lateral cortical surface showed similar BOLD-locked flow dynamics, whereas their cardiac-locked flow differed in both direction and amplitude (Fig. 4C). Together, these examples suggest that cardiac-locked and BOLD-locked CSF signals reflect distinct dynamical processes rather than the same dynamic observed at different timescales, as further supported by brain-wide comparisons described below (Fig. 4D–F).

**Fig. 4.**
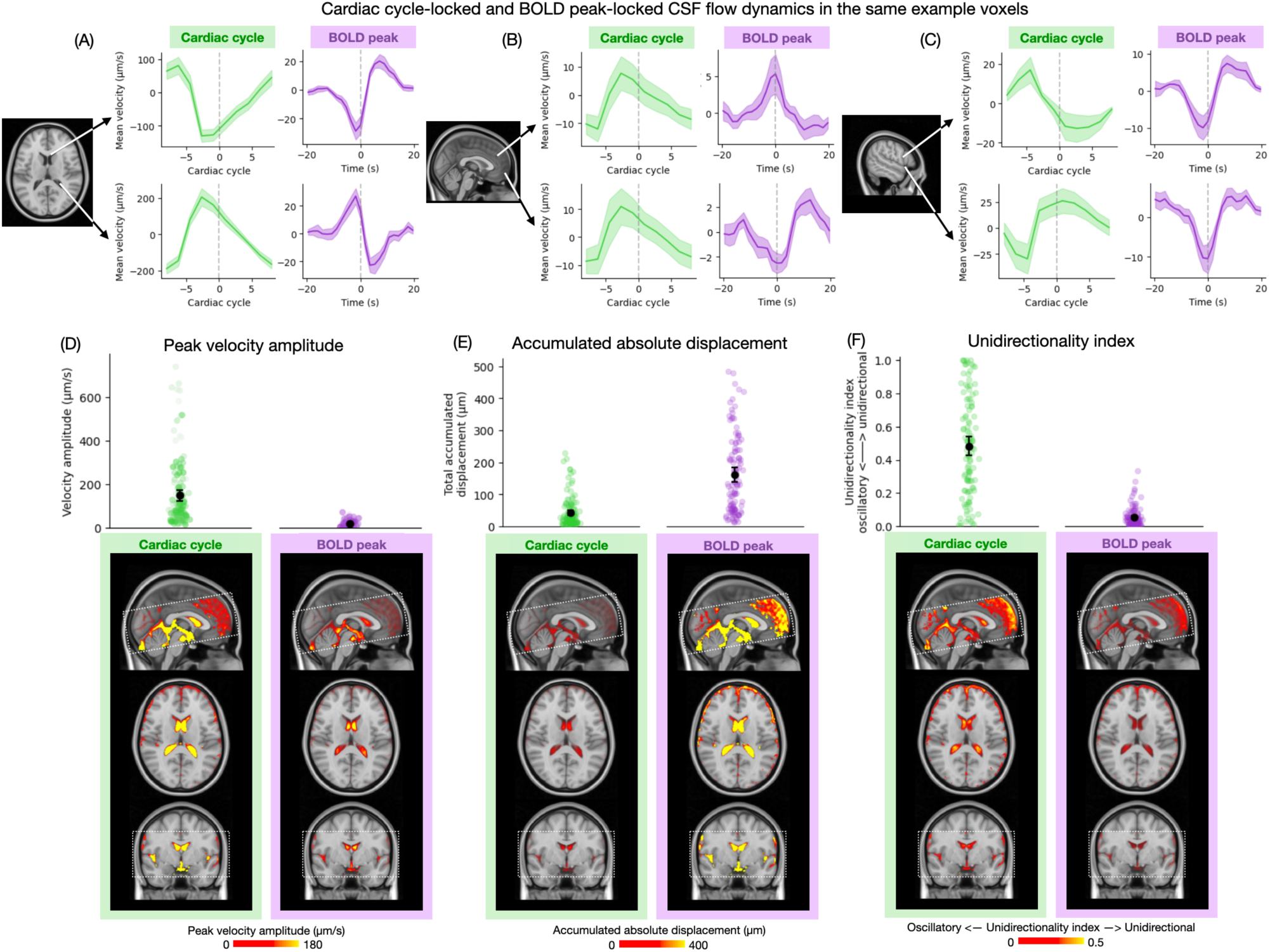
Distinct CSF flow patterns linked to BOLD-oscillations and the cardiac cycle. (A) CSF flow dynamics locked to cardiac cycle and to spontaneous BOLD peaks in the same example voxels (*N* = 17). For each voxel, the left panel shows cardiac-locked CSF velocity, and the right panel shows BOLD-locked CSF velocity. Ventricular voxels located in posterior and anterior portions of the lateral ventricle. (B) Subarachnoid voxels located along the anterior medial wall. (C) Subarachnoid voxels located along the lateral brain wall. Time series are shown as group averages; shaded ribbons indicate standard error across sessions. (D) Velocity amplitude, defined as the peak-to-trough difference of the cycle-locked CSF dynamics. (E) Total accumulated displacement, defined as the time integral of the absolute CSF velocity. (F) Unidirectionality index, defined as the absolute net displacement divided by the total accumulated displacement, where lower values indicate more oscillatory up-and-down motion and higher values indicate more directionally consistent unidirectional flow. In the scatter plots, each dot represents the mean value for one ROI, and black markers with error bars indicate the grand mean ± 95% confidence interval across ROIs. Voxel-wise spatial maps are generated by averaging across subjects. Voxel-wise maps show that cardiac- and BOLD-locked dynamics differ across all three measures, consistent with the ROI-level results. Scatter plots are restricted to subarachnoid ROIs significantly coupled to each cycle; voxel-wise maps include all CSF voxels including the ventricles. Dotted white box indicates acquisition volume placement.

To quantify the differences between the two types of event-locked CSF dynamics, we compared three measures across subarachnoid ROIs significantly coupled to each event type using linear mixed-effects models, with event type (cardiac vs. BOLD) as a fixed effect and a random intercept for ROI (Table S1). We first examined velocity amplitude, defined as the peak-to-trough difference of the cycle-locked CSF dynamics (Fig. 4D). This measure captures the range of the fluid motion during each event, reflecting how strongly CSF is driven back and forth. Cardiac-locked CSF showed substantially greater velocity amplitude than BOLD-locked CSF (Cardiac: mean = 174.64 μm/s, 95% CI [138.89, 210.39]; BOLD: mean = 18.55 μm/s, 95% CI [15.78, 21.32]; LME: β = −155.21, *p* < 0.001), indicating that each cardiac pulse drives much larger instantaneous CSF movement than a spontaneous hemodynamic event.

To capture how much fluid moves over the course of a event, we quantified total accumulated displacement as the time integral of the absolute velocity (Fig. 4E). This measure reflects the total distance a fluid parcel travels during the cycle, regardless of direction, and complements instantaneous velocity by accounting for movement sustained over time. Note that the cardiac cycle unfolds over approximately 1 s, whereas the spontaneous hemodynamic oscillations during sleep unfold over approximately 20 s. Accordingly, despite its smaller instantaneous velocity amplitude, BOLD-locked CSF showed substantially greater accumulated displacement than cardiac-locked CSF (BOLD: mean = 162.07 μm, 95% CI [139.03, 185.12]; Cardiac: mean = 49.43 μm, 95% CI [39.56, 59.30]; LME: β = 116.05, *p* < 0.001). Thus, despite their smaller beat-to-beat velocity, infraslow hemodynamic-driven CSF fluctuations produced substantially greater cumulative absolute movement over the event window than a cardiac cycle.

Finally, to distinguish oscillatory from net-transport flow, we quantified the unidirectionality index (i.e., net-transport fraction), defined as the absolute net displacement (i.e., the magnitude of the *signed* velocity accumulated over a cycle, in which opposite directions cancel) divided by the total accumulated displacement (i.e., the *absolute* velocity accumulated over a cycle, in which all movement accumulates regardless of direction; Fig. 4F). This measure refers to whether flow moves back and forth (i.e., oscillatory flow; low value) or consistently in one direction across the cycle (i.e., unidirectional flow; high value). Cardiac-locked CSF showed a greater unidirectionality index than BOLD-locked CSF (Cardiac: mean = 0.483, 95% CI [0.428, 0.539]; BOLD: mean = 0.055, 95% CI [0.044, 0.066]; LME: β = −0.428, *p* < 0.001). Notably, cardiac-locked dynamics showed a broad range of directional patterns across subarachnoid regions, with some regions showing highly oscillatory responses and others more directionally biased flow, whereas BOLD-locked responses were more uniformly oscillatory overall, consistent with cardiac and hemodynamic drivers producing structurally different modes of CSF motion across the subarachnoid space.

These findings reveal two dissociable modes of CSF motion within the same anatomical spaces. Cardiac pulsation produced brief, high-velocity, more directionally biased flow with strong local heterogeneity; in contrast, BOLD-coupled flow was slower but sustained over much longer time periods, coherent and oscillatory across nearly all subarachnoid regions, and ultimately produced several-fold greater cumulative fluid displacement over the event window.

### Regional dynamics of BOLD–subarachnoid CSF coupling

Infraslow spontaneous BOLD oscillations during sleep reflect neural and hemodynamic activity that is broadly coordinated across cortex, but neurovascular coupling also produces spatially specific responses tied to regional neural activity, generating local tissue expansion and contraction that could displace neighboring CSF (Månsson et al., 2020; Wang, Polimeni, et al., 2026; Wang, Reese, et al., 2026; Zaretskaya et al., 2023). In this scenario of local vascular changes, we would predict that the magnitude of local CSF flow changes would depend on the strength of nearby tissue movement (Fig. 5A). We therefore investigated regional specificity of hemodynamic-driven flow by asking whether BOLD amplitude within a cortical region predicts CSF dynamics in the adjacent subarachnoid space beyond what is explained by global cortical hemodynamic fluctuations.

**Fig. 5.**
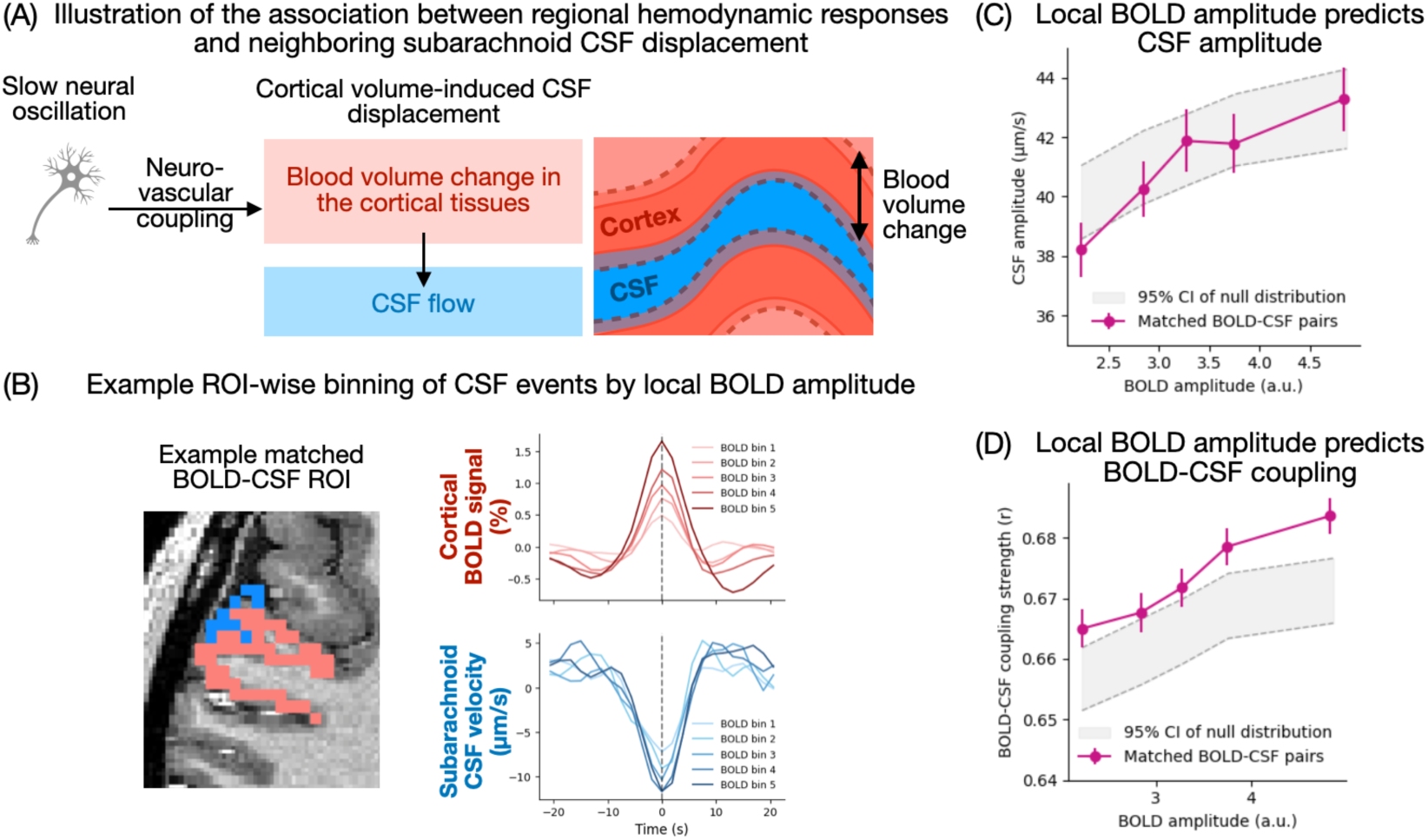
Regional dynamics of tissue-subarachnoid CSF coupling. (A) Schematic of the regional BOLD–CSF pairing. Through neurovascular coupling, sleep-related slow neural oscillations induce local blood volume changes that expand and contract cortical tissue and displace CSF in the adjacent subarachnoid space. Dotted lines around cortical tissue indicate tissue movement, with stronger local movement driving larger local CSF flow changes. (B) Example peak-locked responses from paired tissue–CSF ROI pairs, grouped by local cortical BOLD amplitude. Individual BOLD events were binned by peak-to-trough amplitude, and the same bin assignments were applied to the corresponding paired CSF ROI. (C-D) Quantification of regional BOLD–CSF relationships across paired ROI pairs. Larger local cortical BOLD amplitudes were associated with (C) greater neighboring CSF velocity amplitude and (D) stronger local BOLD–CSF coupling strength, defined as the absolute maximum cross-correlation (*N* = 170 ROI pairs). Magenta lines indicate paired BOLD–CSF pairs. The gray shaded region denotes the 95% confidence interval of the null distribution generated by randomly permuting tissue–CSF pairings, which reflects the BOLD–CSF relationship expected from the shared global signal alone.

Because of the spatial complexity of the subarachnoid space, we defined neighboring tissue– subarachnoid pairs by labeling each subarachnoid voxel with the tissue region it directly borders, so that a matched pair consists of a cortical ROI and the subarachnoid space immediately adjacent to it (Fig. 3A). Within each tissue ROI and subject, we binned individual BOLD-locked events according to the amplitude of the local BOLD oscillation (i.e., peak-to-trough difference in the tissue ROI BOLD signal), and applied the same bin assignments to the corresponding flow dynamics in the paired subarachnoid ROI (Fig. 5B; also see Supplementary Material *Comparison of signal-to-noise ratio across bins*). We then examined whether local BOLD amplitude predicted the amplitude of the CSF velocity change (i.e., peak-to-trough difference in CSF velocity from subarachnoid ROI) as well as the temporal coupling strength between BOLD and CSF changes (i.e., absolute maximum cross-correlation between each tissue–subarachnoid pair; see *Methods*). These measures provided evidence for whether greater local neural and hemodynamic activity predicted stronger, more temporally consistent local CSF dynamics.

We found that cortical regions with larger local BOLD oscillations tended to exhibit both greater CSF velocity change (Fig. 5C, magenta line) and stronger local BOLD–CSF coupling in neighboring subarachnoid regions (Fig. 5D, magenta line). This pattern was confirmed by linear mixed-effects models (see *Methods*), which showed that local BOLD amplitude significantly predicted neighboring CSF amplitude (β = 1.80, *p* < 0.001) as well as BOLD–CSF coupling strength (β = 0.007, *p* < 0.001; Table S2).

To test whether this relationship reflects true spatial specificity rather than a global effect of brain-wide BOLD fluctuations, we randomly permuted the tissue ROI labels and repeated the same analysis using the pseudo-tissue–subarachnoid pairs, to generate a null distribution (1000 shuffles; Fig. 5C & D, gray shaded area). Because this shuffle removes the local correspondence between paired regions but preserves the brain-wide BOLD fluctuation, the null slope reflects the tissue–subarachnoid relationship driven by the global signal. The null distribution itself was centered on a positive slope (Fig. 5C & D, gray shaded area), indicating that global BOLD fluctuations alone could predict some degree of CSF flow response in subarachnoid regions. Relative to this null distribution, local BOLD amplitude predicted both CSF amplitude (permutation test, two-sided *p* < 0.01; Fig. 5C) and BOLD–CSF coupling strength (two-sided *p* < 0.01; Fig. 5D) with significantly steeper slopes than chance, indicating that the relationship between local BOLD amplitude and neighboring CSF dynamics was spatially specific rather than purely driven by shared global fluctuations (also see Table S3; *Comparison of Global and Local BOLD amplitude to predict CSF dynamics* in Supplementary Material for details).

These results indicate that sleep-related hemodynamic oscillations shape CSF dynamics across the subarachnoid space through both mechanisms: a global component consistent with brain-wide blood-volume fluctuations driving coordinated CSF movement, and an additional spatially specific component whereby individual cortical regions with stronger local hemodynamic responses drive greater and more temporally consistent CSF dynamics in their immediately neighboring subarachnoid space, suggesting that regionally specific neurovascular activity during sleep contributes to local fluid displacement beyond global effects.

### Arousal state-dependent CSF flow dynamics modulated by spontaneous hemodynamic responses

Overall, these results so far demonstrate that CSF flow dynamics are associated with spontaneous hemodynamic responses in a coordinated, directional, and spatially specific pattern. We next tested whether this coupled dynamic differs across arousal states. The infraslow hemodynamic oscillations identified as BOLD peaks are characteristic of sleep and have also been observed during drowsiness, but rarely during alert wakefulness (Tagliazucchi & Laufs, 2014); in other words, the BOLD peaks do not typically occur during alertness, preventing comparison of peak-locked flow between sleep and alertness. However, because participants were instructed to keep pressing a button approximately every second for as long as they were awake, our data included sleep periods but also drowsy periods, where participants were not fully alert but still able to press the button but at relatively slow rates. We therefore compared BOLD peaks occurring during periods of behaviorally defined sleep (≤ 8 button presses per 20 s) against those occurring during drowsiness (> 8 presses per 20 s). Only scan sessions with both sleep and drowsy periods were included (*N* = 13 sessions). On average, subjects spent 65.84% of the time asleep (mean 74.19 min, SD = 14.98) and 34.16% drowsy (mean 33.89 min, SD = 9.62). Fewer BOLD peaks were detected during drowsiness than during sleep (drowsy peaks per run: median = 9; sleep: median = 40; Fig. S6; also see *Comparison of signal-to-noise ratio across bins* in Supplementary Material).

We first compared voxel-wise BOLD– CSF coupling strength across arousal states (Fig. 6A). Coupling strength was lower during drowsiness than during sleep in both the subarachnoid space (linear mixed-effects model with arousal state as a fixed effect and scan session as a random intercept: β = −0.067, *p* < 0.001; Table S4) and ventricles (β = −0.056, *p* < 0.01; Table S4), suggesting that the temporal reliability of BOLD–CSF coupling is arousal-state dependent, strengthening during sleep. Because the number of BOLD peaks available for averaging was not matched between arousal states (Fig. S6), we next tested whether this difference in coupling strength persisted when the number of peak events was equated between both states within each run. This analysis was implemented at the subarachnoid ROI level (Fig. 3A) due to the higher signal-to-noise ratio achieved by averaging CSF signal within anatomically defined regions, which was necessary for a trial-matched comparison to be estimable given how few peaks in either state were available in some runs. To achieve matched trial counts, we randomly subsampled peak events from the state with more events in each run, averaging over 20 independent subsamples (see *Methods*). Coupling strength remained higher during sleep than drowsiness after matching trial counts across all ROIs (full sample: β = −0.063, *p* < 0.01; matched sample: β = −0.051, *p* < 0.01; Fig. 6B; Table S4; example subarachnoid ROI in Fig. 6C), indicating that the state-dependent reduction in coupling strength reflects a genuine decrease in the reliability of BOLD–CSF temporal coupling during drowsiness, rather than a mere artifact of sleep contributing more peak events to average than drowsiness. To confirm that the observed patterns described in this work reflect sleep-specific dynamics, we repeated main analyses using sleep-only periods and replicated the key findings (Fig. S7; Table S5). Together, these results suggest that sleep, compared to drowsiness, does not merely provide more opportunities for BOLD peaks to occur, but that the hemodynamic events during sleep are more tightly coupled to CSF movement than those occurring during drowsiness, consistent with the idea that sleep supports more coordinated CSF flow than drowsiness.

**Fig. 6.**
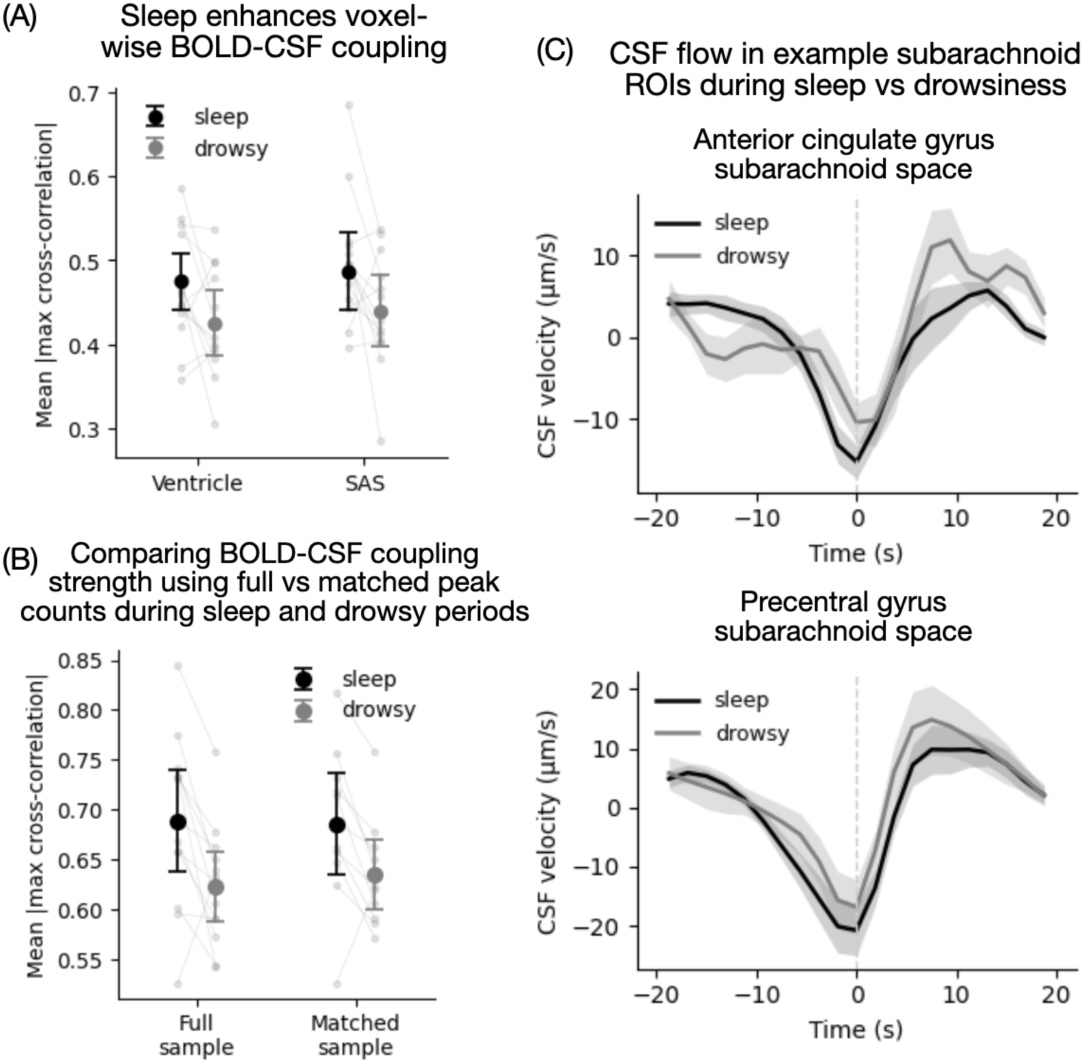
BOLD–CSF dynamics were enhanced during sleep. (A) Group-level BOLD–CSF coupling during sleep vs. drowsiness (*N* = 13 sessions with both sleep and drowsy periods). Voxel-wise absolute maximum cross-correlation between BOLD and CSF flow was calculated and averaged within each CSF compartment (i.e., ventricle, SAS). Coupling strength was lower during drowsiness than during sleep in both compartments. Markers and error bars indicate the group mean 95% confidence interval. Faint lines in the background denote individual subjects. (B) Group-level BOLD–CSF coupling strength using ROI-level analysis, comparing the full sample (all available peaks) and a trial-matched sample (peak events equated between arousal states within each run). Because the number of peak events available for averaging differed substantially between arousal states, we tested whether the reduction in coupling strength during drowsiness was robust to matching trial counts between states within each run. The data show that coupling strength remained significantly lower during drowsiness in both the full and trial-matched samples. Markers and error bars as in (A). (C) Example ROI-wise CSF dynamics during sleep and drowsiness. Subarachnoid BOLD-locked CSF dynamics around precentral gyrus and anterior cingulate gyrus both show more reliably time-locked flow during sleep than drowsiness. CSF velocity is shown separately for sleep and drowsy periods, using all available peak events (full sample). Shaded ribbons indicate SEM across subjects. SAS: subarachnoid space.

## Discussion

By simultaneously imaging brain-wide BOLD activity and CSF velocity during sleep, we found that spontaneous hemodynamic oscillations were coupled to coordinated CSF movement across both the ventricles and subarachnoid space. The findings demonstrate that sleep-related slow CSF dynamics are measurable in the subarachnoid space, extending prior observations of sleep-related CSF flow at the fourth ventricle to a much broader spatial pattern of movement throughout the ventricles and the subarachnoid space. The results further suggest that CSF movement is systematically driven not only by physiological pulsations (∼1 Hz), but also by sleep-related, large-scale hemodynamic events (∼0.05 Hz) that facilitate brain-wide fluid distribution across CSF compartments during sleep (Fultz et al., 2019; Lewis, 2021; Picchioni et al., 2022).

A central advance of this work is the characterization of slow CSF velocity in the human subarachnoid space during sleep. Previous human studies identified large sleep-related CSF inflow signals in the fourth ventricle, but these measurements could not determine how CSF moved across the broader spaces surrounding the cortex. Because the subarachnoid space directly borders the cortical surface, covers surface vessels, and connects with perivascular and cisternal pathways (Rhoton, 2000; Weller et al., 2018; Zhang et al., 1990), sleep-related flow in this compartment is likely to be more directly relevant to fluid exchange between brain tissue and CSF. Solutes that pass from brain tissue into the CSF must then be carried through the subarachnoid space toward efflux sites, including meningeal lymphatics and perineural routes, before leaving the cranium (Aspelund et al., 2015; Louveau et al., 2015; Ma et al., 2017). Our findings show that spontaneous hemodynamic events were coupled to widespread subarachnoid as well as ventricular velocity changes, indicating that sleep-related CSF redistribution was directionally and spatially organized across CSF compartments throughout the brain. Our results show that subarachnoid CSF dynamics were oscillatory around BOLD peaks, with subarachnoid CSF moving largely downward at the BOLD peak, suggesting that sleep-related increases in cortical blood volume underlying the BOLD peak facilitate fluid efflux from the brain. Prior studies inferred this volume exchange from fourth-ventricle inflow signals, which rose as cortical blood volume fell (Fultz et al., 2019; Williams et al., 2023; Zimmermann et al., 2025); our velocity measurements extend this account by directly capturing flow movement in the subarachnoid space. It is also consistent with mouse studies showing that vasodilation promotes solute clearance from the brain: spontaneous and stimulus-evoked vasomotion accelerated paravascular tracer clearance (Van Veluw et al., 2020), increasing blood flow through neurovascular coupling enhanced glymphatic efflux (Holstein-Rønsbo et al., 2023), and slow vasomotion during NREM sleep promoted glymphatic clearance (Hauglund et al., 2025). The precise pathways by which hemodynamic-driven subarachnoid flow exits the brain, and how much of this displaced CSF leaves the cranium rather than redistributing within it, will be an important topic for future investigation.

Fourth ventricle inflow has previously been interpreted as a bulk-flow marker of sleep-related CSF movement (Fultz et al., 2019; Picchioni et al., 2022), but the link between ventricle flow and flow in the broader subarachnoid space has not yet been measured. We found that each fourth-ventricle inflow event was also accompanied by coordinated, largely upward CSF flow throughout both the ventricles and the subarachnoid space, indicating that inflow reflects one component of a much broader flow pattern rather than an isolated ventricular signal. The spatial pattern of these inflow-driven CSF changes resembles the hemodynamic-driven pattern but with largely inverted direction, indicating that the two measurements captured the same brain-wide CSF cycle at different points in time. Together, the combined hemodynamics, ventricular inflow, and subarachnoid dynamics identify a coherent, continuous pattern: as cortical blood volume falls following a hemodynamic peak, CSF enters the fourth ventricle from the cervical subarachnoid space (i.e., the inflow peak), while CSF across most of the subarachnoid space flows consistently upward toward the dorsal convexity. Approximately 7 seconds later, as the next cortical hemodynamic response peaks and blood volume increases, the pattern reverses and most subarachnoid CSF is displaced downward and CSF flows downward through the fourth ventricle (Fig. S8). This alternating pattern of CSF motion, driven by continuous infraslow hemodynamic oscillations, is consistent with a sleep-specific mechanism that could facilitate solute transport and clearance.

Importantly, these hemodynamic-driven flow dynamics are distinct from the cardiac-driven dynamics. Cardiac pulsation generated large, transient velocity changes and, in many regions, displaced CSF preferentially in one direction during each event, likely reflecting rapid pressure-wave propagation through anatomically constrained pathways (see Wang, Reese, et al., 2026). In contrast, sleep-related infraslow hemodynamic-driven flow was more oscillatory and, despite its smaller instantaneous velocity, produced substantially greater cumulative displacement due to its longer duration. Because a single hemodynamic event spans tens of cardiac cycles, this reflects the size of each slow excursion, rather than total motion per unit time. The two drives therefore appear to move CSF in different ways: cardiac pulsation produces brief, directionally biased flow that, repeated with each beat, may contribute to net transport along specific pathways, whereas slow hemodynamic oscillations during sleep produce large, spatially coordinated oscillatory displacements that may promote fluid mixing and exchange across CSF compartments.

The brain-wide, hemodynamic-driven CSF flow pattern was spatially organized, but not uniform. Most subarachnoid regions, concentrated along the dorsal cortical surface, showed downward CSF flow at the BOLD peak and upward flow at the inflow peak, with similar peak velocity amplitude at both events, consistent with cortical-adjacent CSF being displaced as cortical blood volume rises and drawn back as it falls. In contrast, subarachnoid regions in the basal cisterns and basal cortex maintained the same upward flow direction at both BOLD and inflow peaks and were much more strongly coupled to fourth-ventricle inflow than to cortical BOLD, consistent with their direct continuity with the fourth ventricle and spinal subarachnoid space. Because this flow ran opposite to the downward direction of net drainage throughout the event, sleep-related oscillations in these central compartments may promote CSF circulation and mixing rather than simple outflow. This anatomical organization suggests that sleep-related CSF flow is shaped by multiple interacting components: a widespread component at the brain surface that tracks cortical blood volume, and an inflow-dominated component near central CSF compartments that is tied to bulk exchange with the spinal subarachnoid space.

The spatial heterogeneity in flow patterns described above suggests that local CSF flow is shaped by neighboring brain tissue that expands and contracts with regional neural and hemodynamic activity through neurovascular coupling. Animal studies have linked perivascular CSF flow to these local vascular dynamics at the microscale (Bojarskaite et al., 2023; Kedarasetti et al., 2020; Van Veluw et al., 2020), and recent human work has shown that CSF mobility can be driven locally, with 0.1-Hz visual stimulation increasing CSF mobility in visual cortex (Hirschler et al., 2025). However, these measures index the magnitude of local CSF motion rather than its direction, leaving open how spontaneous hemodynamic events organize the direction of flow across compartments. We found that local cortical BOLD amplitude predicted CSF dynamics in adjacent subarachnoid regions, indicating that regional neurovascular activity directly shapes nearby CSF flow. This result is consistent with mechanistic models from animal studies and suggests that the underlying sleep-related neural activity shapes CSF movement in a regionally specific manner, potentially contributing to spatial differences in fluid exchange rate and protein clearance efficiency across brain regions (Han et al., 2023; Harrison et al., 2020).

Several limitations should be considered. First, the dataset encoded velocity along the head–foot direction only, the direction with the major flow component, so the main conclusions about directional flow are axis-specific. Second, temporal resolution likely limited the sensitivity to detect rapid fluid propagation delays. Although the observed time-locked patterns across CSF compartments could reflect a temporal sequence of flow propagation, the TR value (1.78 s) was likely too coarse to resolve such fine temporal offsets, and the lag-at-maximum-correlation maps did not reveal strong spatial variation. Third, our low VENC was optimized for slow subarachnoid flow, and therefore faster ventricular flow events may be partially attenuated by phase-wrap exclusion. Fourth, although the regional pairing analyses supported local tissue–CSF relationships, partial-volume effects and shared boundary-related MR signal could contribute to spatial specificity. Lastly, sleep was inferred from behavioral responsiveness alone. As a result, periods classified as sleep should be interpreted as behaviorally defined reduced-arousal periods. We therefore cannot determine sleep stage or fully exclude quiet wakefulness or drowsiness.

In summary, we show that during sleep, global and local infraslow hemodynamics coordinate CSF redistribution, producing spatially organized, directional flow patterns throughout both ventricular and subarachnoid compartments. This work establishes a framework for understanding how large-scale brain oscillations shape human CSF dynamics during sleep and provides a foundation for determining how these coordinated fluid movements contribute to metabolite exchange, waste clearance, and brain fluid homeostasis (Eide et al., 2021; Iliff et al., 2012; Xie et al., 2013).

## Methods

### Participants

Subjects were screened not to have any neurological, psychiatric, or sleep disorders and not be currently taking psychiatric or sleep medications. Subjects were asked to avoid caffeine for 24 hours prior to the study. All subjects provided written informed consent. Data were collected across 20 scanning sessions from 11 subjects. To maximize the amount of sleep data collected, subjects who were able to sleep in the scanner were invited to return for up to two additional sessions: four subjects completed three scanning sessions, one completed two, and six completed one. A scanning session refers to a single overnight scan visit, consisting of one to four 45-minute functional runs; repeated sessions from the same subject were acquired on different nights and treated as separate observations in all analyses. After motion exclusion (see *fMRI and CSF-flow preprocessing*), 17 sessions from 9 subjects (8 female; age: mean = 26.1 years, SD = 5.0 years) were retained for analysis. All experimental procedures were approved by the Massachusetts General Hospital Institutional Review Board.

### MRI acquisition

Participants were scanned supine in a 7T Siemens MAGNETOM Terra scanner (Siemens Healthineers, Erlangen, Germany) using a 32-channel head coil (Nova Medical, Wilmington, MA, USA). A T_1_-weighted anatomical multi-echo MPRAGE scan was first acquired as an anatomical reference at 0.75 mm isotropic resolution. Scan start time was scheduled according to each participant’s habitual bedtime (between 10:00 PM and midnight). Each session consisted of 1–4 45-min functional runs, during which participants were instructed to alternately press a button with their index and middle fingers approximately 1 s apart and to resume the task after spontaneous awakenings. Arousal state was inferred behaviorally from button-press performance: 20-s windows with ≤ 8 presses were classified as sleep and those with > 8 presses as drowsy.

BOLD and CSF-flow data were acquired simultaneously using SOPHI-EPTI sequence at 7 Tesla (Dong et al., 2023; Wang et al., 2024; Wang, Polimeni, et al., 2026; Wang, Reese, et al., 2026) with echo-planar time-resolved imaging (Dong et al., 2024; Wang et al., 2019). EPTI acquires a train of echoes after each excitation, so that a single readout yields distortion-free images across a range of echo times rather than at the single echo available in conventional phase-contrast acquisitions. Velocity encoding uses a pulsed-gradient spin-echo (PGSE) scheme whose extended encoding time achieves a low VENC without substantial diffusion-induced CSF signal loss, and the asymmetric spin echoes resolved within the EPTI readout provide T2*-weighted contrast for BOLD measurement (Wang, Polimeni, et al., 2026; Wang, Reese, et al., 2026). From the same acquisition, the reconstructed magnitude images therefore provided the BOLD-sensitive signal, while the phase images provided quantitative velocity-encoded CSF-flow measurements. The experiment used single-direction velocity encoding along the foot–head (*z*) axis, with 26 oblique slices and TR = 1780 ms. The acquisition slab was positioned to include the lateral and third ventricles and the mid-axial subarachnoid space, with its bottom slice positioned to capture CSF inflow at the fourth ventricle (Fig. 1A). All scans were acquired at 2-mm isotropic resolution (FOV = 216 × 192 mm²; matrix = 108 × 96; multiband factor = 2) with a 68-echo EPTI readout. The EPTI readout sampled signal across 68 echo times (TE range: 63.4–112.6 ms) and was centered on a spin-echo refocusing time of 88 ms. A VENC of 1.6 mm/s (effective b ≈ 80 s/mm²) was chosen to capture slow CSF flow in the subarachnoid space. Each run began with 20 non-velocity-encoded volumes, which were later used as a phase reference for CSF-flow estimation (Dong et al., 2023, 2025; Wang et al., 2024; Wang, Polimeni, et al., 2026).

The cardiac and respiratory cycles were recorded simultaneously during scanning. Cardiac pulse was measured using a piezoelectric pulse transducer placed on the left thumb (AD Instruments, Colorado Springs, CO, USA), and respiration was measured using a piezoelectric respiratory belt placed around the chest (UFI, Morro Bay, CA, USA). Both signals were recorded using LabChart version 7. The cardiac signal was low-pass filtered at 10 Hz, and the respiratory signal was band-pass filtered between 0.005 and 0.8 Hz, and cardiac and respiratory cycles were identified from peaks in the filtered signals.

### fMRI and CSF-flow preprocessing

Raw EPTI data were reconstructed into multi-echo complex-valued images. BOLD signal was estimated from magnitude-valued images spanning TE = 68.5–106.9 ms. Because spin-echo contrast is predominantly T2-weighted and has limited BOLD sensitivity, we used this range, which includes asymmetric spin echoes on both sides of the refocusing time, providing T2*-weighted contrast (Wang et al., 2021; Wang, Polimeni, et al., 2026). The velocity-encoding gradients imparted mild diffusion weighting to the magnitude images, which reduces SNR and attenuates intravascular signal but leaves the contrast BOLD-dominated at this b-value (∼ 80 s/mm²). Rigid-body motion parameters were estimated from the magnitude images using AFNI (Cox, 1996). The transformations were applied to each echo’s magnitude image (cubic interpolation) and to the phase image (nearest-neighbor interpolation). The BOLD image was then computed as the mean of the motion-corrected magnitude images across echoes to increase signal-to-noise ratio. Slice acquisition timing was computed and tracked through the motion correction step using nearest-neighbor interpolation, preserving per-voxel timing. Slice timing correction was applied to the BOLD images to align all slices to a common reference time within each TR. Each run’s BOLD image was registered to subject’s T_1_-weighted anatomical scan.

Flow velocity was calculated from echo-averaged phase images spanning TE = 80.5–94.9 ms, the central echoes near the spin-echo refocusing time, which are less sensitive to confounding phase fluctuations. Because CSF has a longer T_2_ relaxation time than tissue or blood, especially at 7T, its signal is selectively preserved at these longer echo times, making this long-TE image more specific to CSF flow. In addition, the velocity-encoding gradients suppress residual blood signal through intravoxel dephasing of rapidly flowing spins. To remove spatially smooth phase variations, we used a third-order spatial polynomial fit to non-CSF voxels on each slice and time frame and subtracted it from all voxels. Motion correction transformations were applied as described above. For background phase correction, the first 20 non-velocity-encoded volumes were averaged to form a phase reference. Phase differences between velocity-encoded volumes and this reference were computed voxel-wise. A second third-order spatial polynomial fit was applied to remove residual phase offsets from eddy currents, and the corrected phase difference was converted to velocity (Wang, Polimeni, et al., 2026). Example voxel time series for BOLD and flow estimation can be found in Fig. S1. All analyses were performed in each subject’s motion-corrected functional space, hereafter native functional space.

Within-ROI spatial smoothing (FWHM = 2.5 mm) was applied separately within each mask (CSF, cortical gray matter, all gray matter, and white matter; see *CSF, subarachnoid space, ventricle, gray matter, and white matter masks*) to prevent spatial blurring across ROI boundaries. ROI-specific time series for both brain tissue and CSF spaces were extracted within each smoothed mask.

Runs were excluded if ≥ 20% of volumes exceeded 0.3 mm framewise displacement (FD); in total, 14 runs were excluded due to motion. Volumes with large motion (> 0.3 mm) were excluded from all analyses. On average, in the remaining 43 runs, 2.14% (SEM = 0.32%) of volumes per run were excluded due to motion. To reduce contamination from phase wrapping, voxels with absolute velocity exceeding 90% of the VENC (1.44 mm/s) in more than 5% of time points were excluded from CSF flow analysis. On average, 2.75% (SD = 1.78%) of voxels per run were excluded due to phase wrapping. For analyses focused on spontaneous BOLD-associated CSF dynamics, slice-timing-specific cardiac variance was regressed from the flow velocity time series to reduce cardiac noise.

### CSF, subarachnoid space, ventricle, gray matter, and white matter masks

Cortical surface reconstruction and parcellation were performed automatically using FreeSurfer’s recon-all pipeline (Fischl, 2012) applied to each subject’s T_1_-weighted MPRAGE, producing a whole-brain segmentation (i.e., aparc+aseg) in which each voxel is assigned to a specific brain tissue region or ventricle. To identify all CSF voxels, including the subarachnoid space, while minimizing contamination from adjacent brain tissue, we generated run-specific masks using a multi-step procedure. First, because CSF appears relatively bright in long-TE EPTI images, we computed the time-averaged functional image for each scan and selected voxels with mean intensity above the 85th percentile as an initial CSF inclusion mask. Second, to identify non-CSF tissue, we transformed the brain tissue mask (derived from aparc+aseg) into functional space using trilinear interpolation, yielding a tissue overlap map. Voxels where the tissue overlap probability was below 0.2 were retained as likely CSF voxels. Third, as a safeguard, we excluded any remaining voxels labeled in aparc+aseg as gray matter or white matter. The final CSF mask was defined as the intersection of the high-intensity functional mask and the high-likelihood CSF overlap mask, and the removal of all identified brain-tissue voxels.

Gray and white matter masks were defined from the aparc+aseg parcellation, after excluding any voxels in the CSF mask described above. Gray matter was further subdivided into a cortical gray matter mask (using FreeSurfer cortical labels) and a subcortical gray matter mask (using subcortical structure labels). The ventricle mask was identified using the ventricle labels from the aparc+aseg parcellation, which were dilated by one voxel to ensure complete spatial coverage of the ventricle boundary, then intersected with the full CSF mask. The subarachnoid space mask was defined as voxels within the CSF mask excluding those identified as ventricles. Subarachnoid space ROI labels were assigned based on the nearest neighboring tissue labels, with each voxel labeled according to the majority label among its closest voxels in the brain tissue parcellation. All masks were obtained in subject’s run-level native functional space.

### Fourth ventricle inflow voxel identification

Fourth ventricle inflow voxels were identified from the magnitude images based on their signal intensity, then spatially restricted to the fourth ventricle mask. Because inflow-related signal is expected to appear as high-intensity transients relative to surrounding tissue (Fultz et al., 2019; Williams et al., 2023), voxels with intensities exceeding the 95th percentile of all voxels in the first volume were first identified as putative inflow voxels. The fourth ventricle mask was then dilated by one voxel to ensure complete spatial coverage of the ventricle boundary, and the intersection of this dilated mask and the high-intensity voxels within the bottom three acquired slices was defined as the inflow ROI. One scanning session (comprising one single run) was excluded because no voxels survived the intensity threshold, resulting in a final sample size of *N* = 16 for inflow-related analyses.

### BOLD and fourth ventricle inflow peak identification

BOLD and fourth ventricle inflow peaks were identified from magnitude images and subsequently used to time-lock both BOLD-like magnitude and phase-derived flow images. Global BOLD peaks were identified from the cortical gray matter mean magnitude time series (cortical gray matter voxels were defined using FreeSurfer cortical labels from the aparc+aseg parcellation; see *CSF, subarachnoid space, ventricle, gray matter, and white matter masks*). Signals were converted to percent signal change and band-pass filtered to capture infraslow oscillations during sleep (0.01–0.15 Hz). Peaks were identified with a minimum height above the 50th percentile of the time series, a minimum prominence equal to half the difference between the 99th and 1st percentiles, and a minimum interpeak distance of 10 s (Fig. 1B). Peaks occurring during or within ±5 s of frames with framewise displacement >0.3 mm were excluded to minimize motion-induced artifacts.

Fourth ventricle inflow peaks were identified from the mean magnitude time series extracted from the fourth ventricle inflow mask defined above. The fourth ventricle inflow time series was low-pass filtered (0.15 Hz). Peaks were identified using the same height, prominence, and minimum-distance criteria as for BOLD peaks. Given the susceptibility of inflow signals to motion-related artifacts, peaks occurring during or within ±10 s of frames with framewise displacement >0.3 mm were excluded.

### MNI space normalization

All computations and statistical analyses were performed in subject’s native functional space; MNI normalization was used only to visualize group-level results. Subject-level maps were transformed from native functional space to MNI152 standard space using ANTs (Avants et al., 2011), by applying the functional-to-anatomical and anatomical-to-MNI transformations using nearest-neighbor interpolation for binary masks and trilinear interpolation for continuous-valued maps. To create a group-level CSF masks, we counted, for each MNI voxel, the number of sessions in which it was labeled CSF and retained voxels labeled CSF in more than 3 of the 17 sessions; clusters smaller than 10 voxels were removed. Continuous-valued maps were temporally smoothed (2-point moving average) and averaged across sessions, and the group-level CSF mask was applied to restrict visualization to CSF voxels.

### Significance testing of BOLD–CSF coupling

To assess whether CSF velocity changes were temporally coupled to the BOLD signal, we performed a voxel-wise cross-correlation analysis between CSF and global cortical BOLD using time-locked data. For each subject and run, we first computed the mean cortical BOLD time course by averaging the cortical BOLD signal over a window extending 20 s before and after each peak event within a run (i.e., overall BOLD signal in all cortical voxels). For each CSF voxel, we extracted the corresponding time-locked CSF velocity time series within the same window (i.e., CSF signal in each CSF voxel). Both the BOLD and CSF time series were z-scored within the ±20 s time window, and the CSF data were additionally smoothed with a 2-point moving average, as CSF velocity changes driven by slow hemodynamic events are expected to vary smoothly over time. For each CSF voxel, we computed the normalized cross-correlation between the voxel-wise CSF and mean BOLD time series across lags ±20 s from event peaks and used the maximum absolute correlation as the voxel-wise test statistic for BOLD–CSF temporal coupling.

To test whether the observed coupling exceeded what would be expected from temporal autocorrelation and spectral properties of the CSF signal alone, we constructed a null distribution using phase randomization of signals for each CSF voxel (1000 shuffles). For each permutation, we randomized the Fourier phases of the voxel’s time-locked CSF signal while preserving its amplitude spectrum, then applied the inverse transform to generate a shuffled CSF signal. This procedure preserves the power spectrum and autocorrelation structure of the CSF signal but destroys its specific temporal phase relationships. For each phase-randomized CSF signal, we repeated the cross-correlation between the shuffled CSF signal and the same mean BOLD signal to generate a null distribution for each voxel using the maximum absolute correlation from each iteration. A p-value for each voxel was calculated as the proportion of permuted cross-correlations exceeding the observed value. The significance testing of BOLD–CSF coupling in the subarachnoid–tissue parcellation (i.e., ROI-based analysis) was performed using the same procedure. Inflow–CSF coupling, at both the voxel and ROI level, was assessed using the same procedure, with the time-locked fourth-ventricle inflow signal as the reference.

### CSF dynamics metrics

We used multiple measures to characterize CSF dynamics around cortical BOLD peaks, fourth-ventricle inflow peaks, and cardiac cycles. These metrics capture distinct properties of the flow: (i) *Velocity amplitude,* computed as the peak-to-trough difference of the event-locked CSF velocity waveform, quantified the size of the velocity range during each event. (ii) *Coupling strength*, computed as the maximum absolute cross-correlation between voxel-wise CSF velocity and the reference waveform, quantified how reliably CSF velocity followed the timing of the reference signal. (iii) *Signed coupling*, computed as the cross-correlation at the lag of maximum absolute correlation, captured whether the flow deflection was positive-going or negative-going relative to the reference event. (iv) Absolute peak velocity, computed as the maximum absolute velocity within the event-locked window, was used to compare coupling groups. For the cardiac-versus-BOLD comparison, we computed two additional measures: (v) *Total accumulated displacement*, computed as the time integral of the absolute velocity over the cycle-locked window, captured the total distance a fluid parcel travels regardless of direction; and (vi) *Unidirectionality index*, computed as the absolute net displacement divided by the total accumulated displacement, quantified whether flow reciprocated back and forth (values near 0) or moved consistently in one direction (values near 1). All three cardiac-versus-BOLD comparisons (velocity amplitude, total accumulated displacement, and unidirectionality index) remained significant after Bonferroni correction for three comparisons (corrected p < 0.001).

### Spatial specificity linear mixed-effects models and permutation test

Linear mixed-effects models were used to test whether CSF dynamics varied as a function of BOLD amplitude. Two CSF outcomes were examined: (i) CSF amplitude, defined as the peak-to-trough difference over the time-locked cycle, and (ii) CSF– BOLD coupling strength, defined as the absolute maximum cross-correlation between CSF and BOLD time courses. We used ROI-based data derived from aparc+aseg parcellations for tissue signals, together with the corresponding subarachnoid space ROI definition (see above) for CSF signals. Because CSF parcellation was defined using its surrounding tissue labels, CSF–tissue pairs with the same label names are by definition spatially adjacent, and unmatched pairs are more likely to be further in distance.

For tissue signals, BOLD amplitude was defined as the peak-to-trough difference over the time-locked cycle. Trial-level time-locked BOLD responses were sorted into five equal-frequency bins according to BOLD amplitude, ranging from low to high amplitude. Using these BOLD-defined bins, we then computed the corresponding CSF amplitude and CSF–BOLD coupling strength within each bin. We fit two linear mixed-effects models, using BOLD amplitude as a fixed effect and each CSF outcome as the dependent variable. Models included scan session as a random intercept and ROI as an additional random intercept via a variance-components term, treating ROI-to-ROI variability as a nuisance source of variance.

For the permutation test of spatial specificity, we refit the same linear mixed-effects model for each permutation in which CSF ROI labels were randomly scrambled, generating a null distribution based on mismatched BOLD–CSF ROI pairs. The observed (true, unscrambled) β was then compared with this null distribution, and a permutation *p* value was computed as the proportion of null β values at least as extreme as the observed β (two-tailed). Significant results (*p* < 0.05) indicate spatial specificity, such that anatomically adjacent BOLD–CSF ROI pairs exhibit a stronger BOLD-amplitude dependence than expected under spatial scrambling.

### Trial-matched ROI-level comparison of arousal-state effects on BOLD–CSF coupling

Because the number of usable BOLD peaks differed substantially between sleep and drowsiness within a given run (Fig. S6), we performed an additional analysis to test whether the reduction in coupling strength during drowsiness reflected a true state-dependent difference rather than the disparity in the number of peak events available for trial-averaging. For each subarachnoid ROI (Fig. 3A) and each run in which both sleep and drowsy peaks were present, we compared coupling strength for two conditions: (i) a full-sample condition using all available peak events for each state, and (ii) a trial-matched condition in which, for each run, the arousal state with more available peak events was subsampled without replacement to equal the number of events available in the other state (drowsy peaks per run: median = 9, range = 1–63; sleep peaks per run: median = 40, range = 1– 62). The full-sample condition used all 13 sessions with both sleep and drowsy periods; the trial-matched condition used 11 sessions, as 2 of the 13 sessions did not have more than one peak event in both arousal periods to support trial-matching.

To reduce sensitivity to any single random subsample, the trial-matched trace was computed by averaging 20 independent random subsamples of the appropriate size prior to cross-correlation. For both conditions, the trial-averaged CSF signal within each ROI was cross-correlated against the run’s cortical BOLD reference and tested for significance using the same phase-randomization procedure described above (Significance testing of BOLD–CSF coupling). Linear mixed-effects models were estimated with arousal state as a fixed effect and scan sessions as a random intercept, fit separately for the full-sample and trial-matched conditions (Table S4).

## Acknowledgments

This work was supported by National Institutes of Health grants U19NS128613, R01AT011429, R01AG070135, R00AG083056, R01EB036507, and U24NS129893, the McKnight Scholar Award, the Sloan Fellowship, the Pew Biomedical Scholar Award, and the Simons Collaboration on Plasticity in the Aging Brain (811231). This work also used instrumentation supported by the NIH Shared Instrumentation Grant Program (S10-OD023637). We thank Banban Tan, Nicholas Cicero, Daniel Gomez, Ewa Beldzik, Maikala Banks, Baarbod Ashenagar, Harrison Fisher, Brandon Dormes, and Courtney Zambello for their help with subject recruitment and data acquisition.

