## Supplemental material for "Spontaneous Hemodynamic Oscillations during Sleep Coordinate Directional CSF Flow in the Subarachnoid Space and Ventricles"

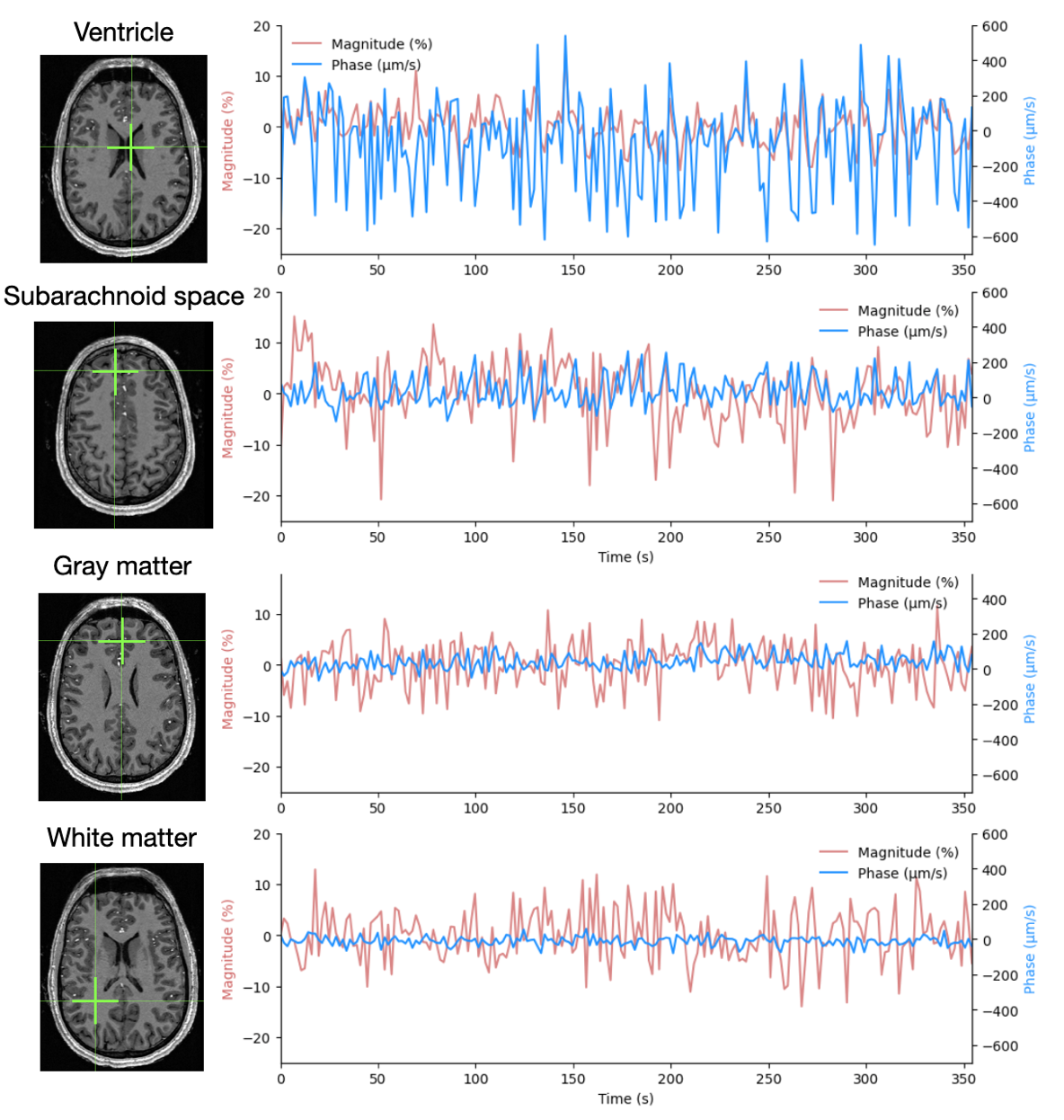

**Fig. S1. Example preprocessed magnitude and phase time series from voxels in different brain compartments.** Traces show segments of the preprocessed magnitude and phase signals from an example subject. Magnitude signals in red reflect BOLD-like signals. Phase signals in blue reflect velocity signals.

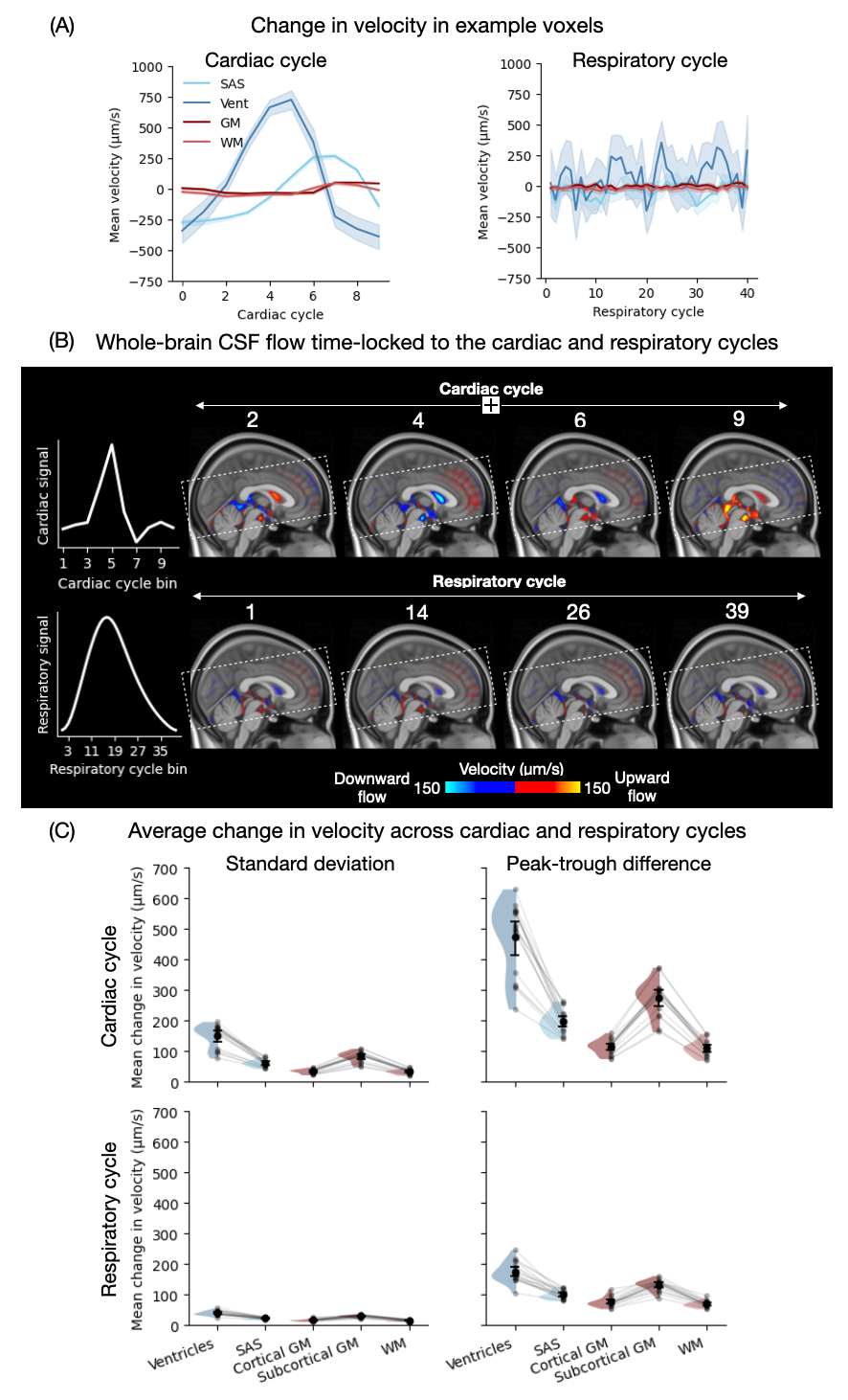

**Fig. S2. CSF flow velocity modulated by cardiac and respiratory cycles.** (A) Example voxel-wise cycle-locked CSF flow. Example voxels in the ventricles and subarachnoid space exhibited strong cardiac-locked CSF flow dynamics, whereas respiratory-locked effects were weaker. (B) Whole-brain cycle-locked CSF flow patterns. Whole-brain averages in standard MNI space revealed robust cardiac-locked flow changes with rapid alternation in flow direction, while respiratory-locked effects were smaller. (C) Quantification of cycle-locked velocity changes across CSF and tissue compartments. Standard deviation and peak-to-trough difference showed the largest velocity changes in ventricular CSF flow, followed by subcortical gray matter tissue motion, subarachnoid space CSF flow, cortical gray matter tissue motion, and white matter tissue motion. The same rank order was observed for both cardiac and respiratory cycles, with weaker effects for respiration.

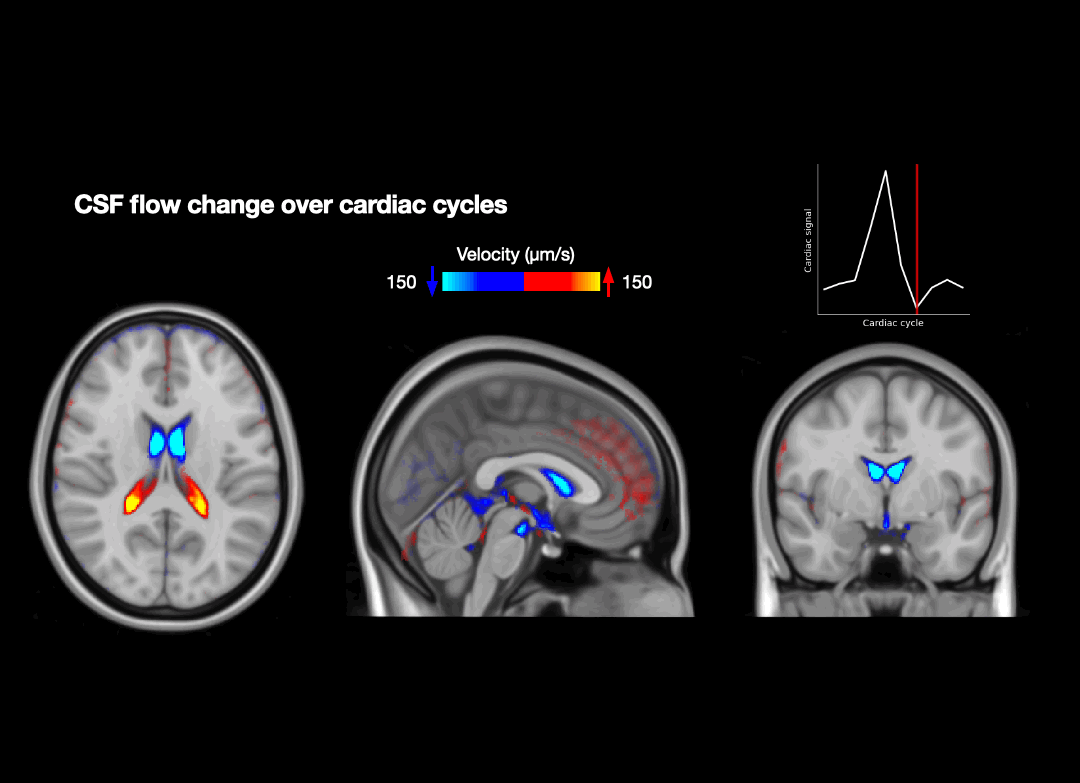

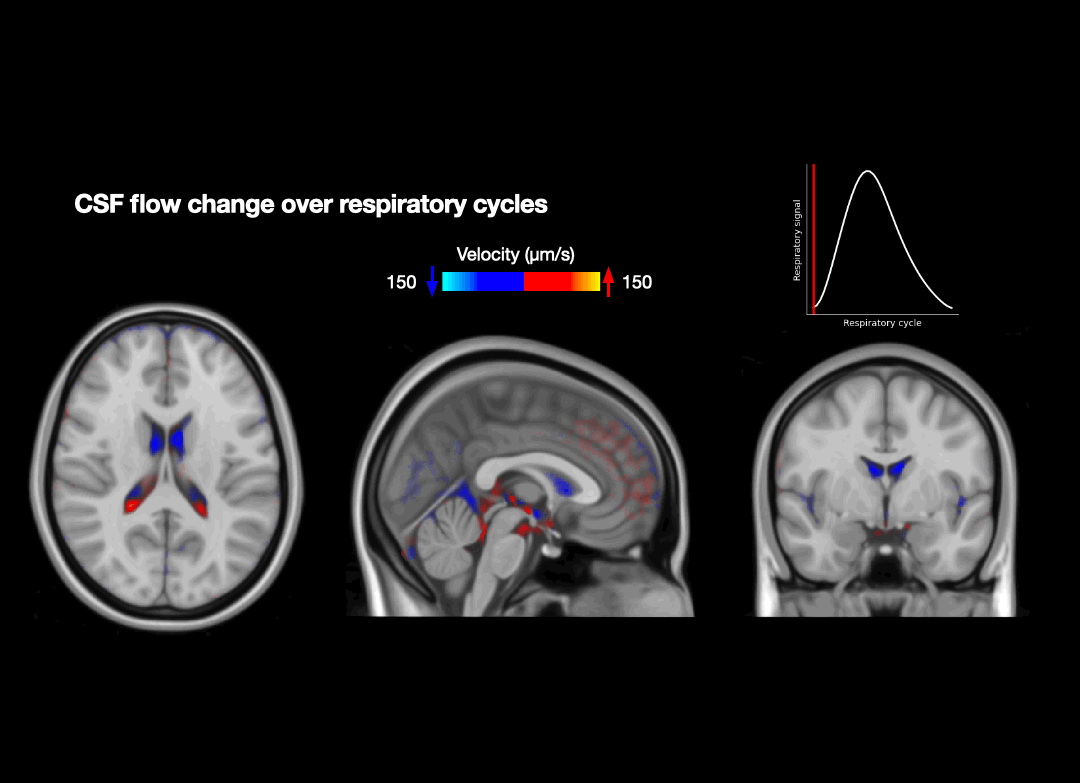

**Movie S1.** Whole-brain cardiac cycle- and respiratory cycle-locked CSF flow patterns. Playback is accelerated to a total duration of 1 second for the cardiac cycle and 4 seconds for the respiratory cycle.

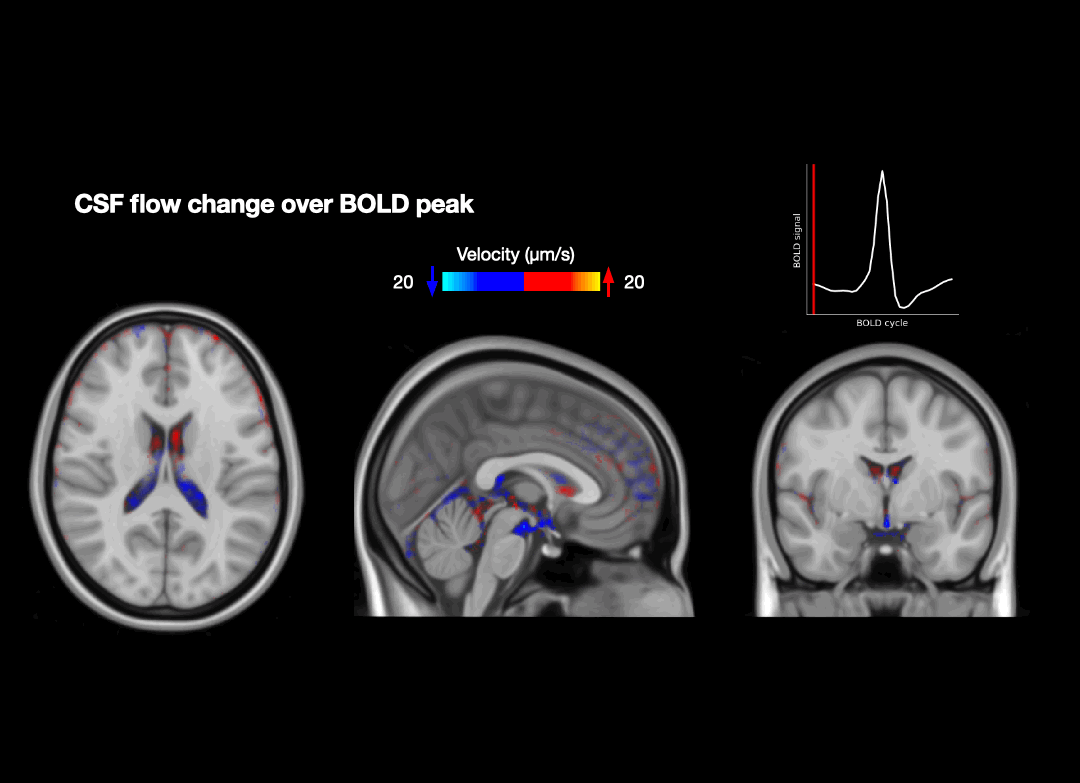

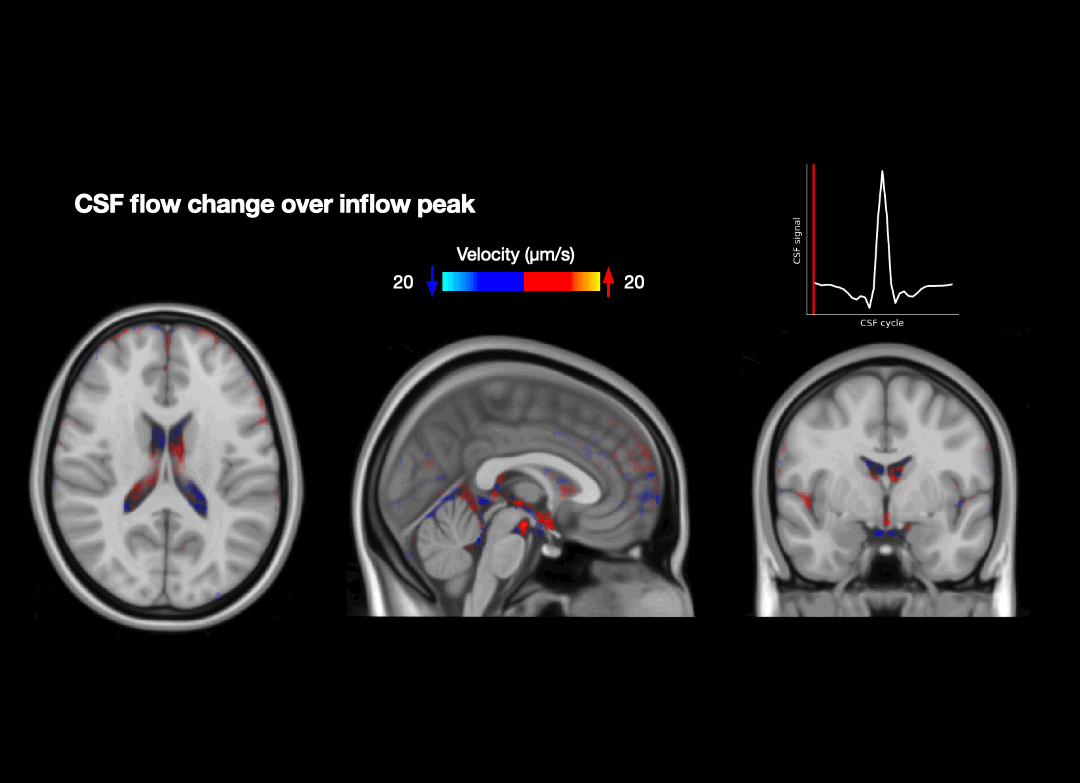

**Movie S2.** Whole-brain BOLD-locked and inflow-locked CSF flow patterns. Playback is accelerated to a total duration of 5 seconds.

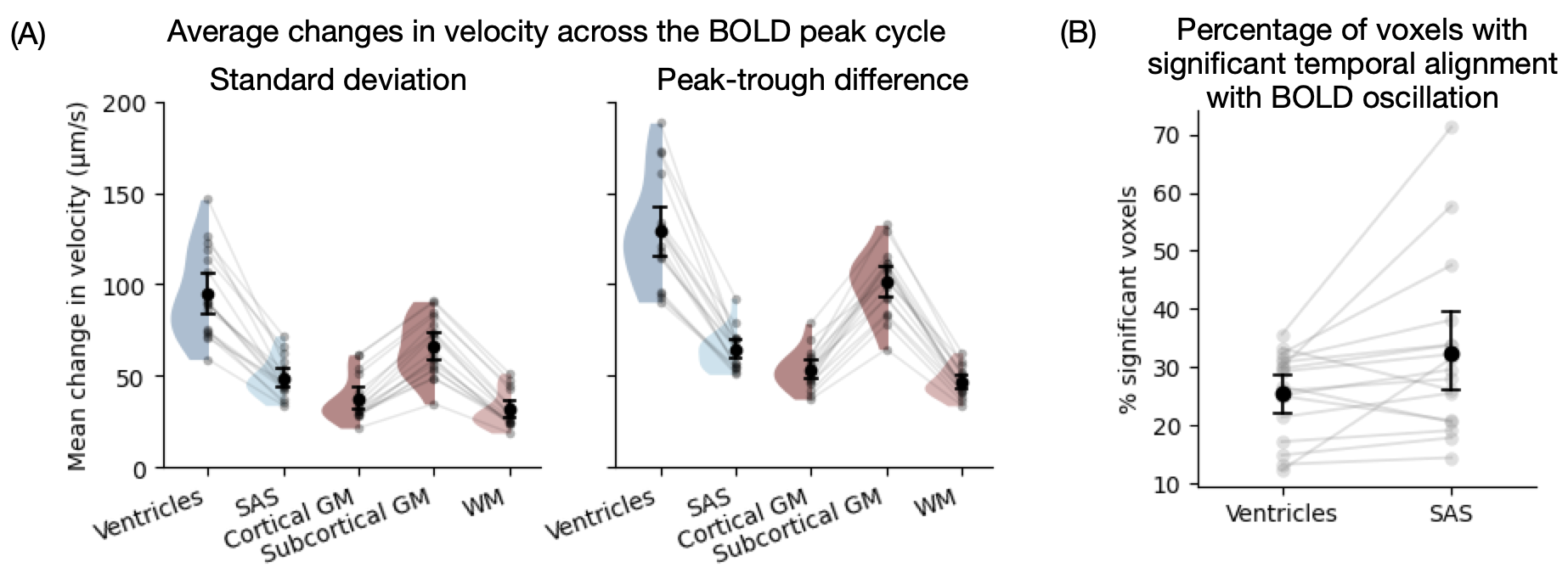

**Fig. S3. CSF flow velocity modulated by spontaneous hemodynamic oscillations.** (A) Average changes in velocity across BOLD peak cycle, for each tissue type. The standard deviation and peak-to-trough difference were calculated for each voxel and averaged within ventricle, subarachnoid space, cortical gray matter, subcortical gray matter, and white matter masks. Gray lines and dots indicate subject averages. All pairwise differences between ROIs were significant (paired *t*-tests, *p* < 0.01). (B) Percentage of voxels showing significant (*p* < 0.05) temporal coupling with BOLD oscillations within subarachnoid space and ventricle masks. Dots indicate subject averages. Group level averages in black; error bars are 95% CI.

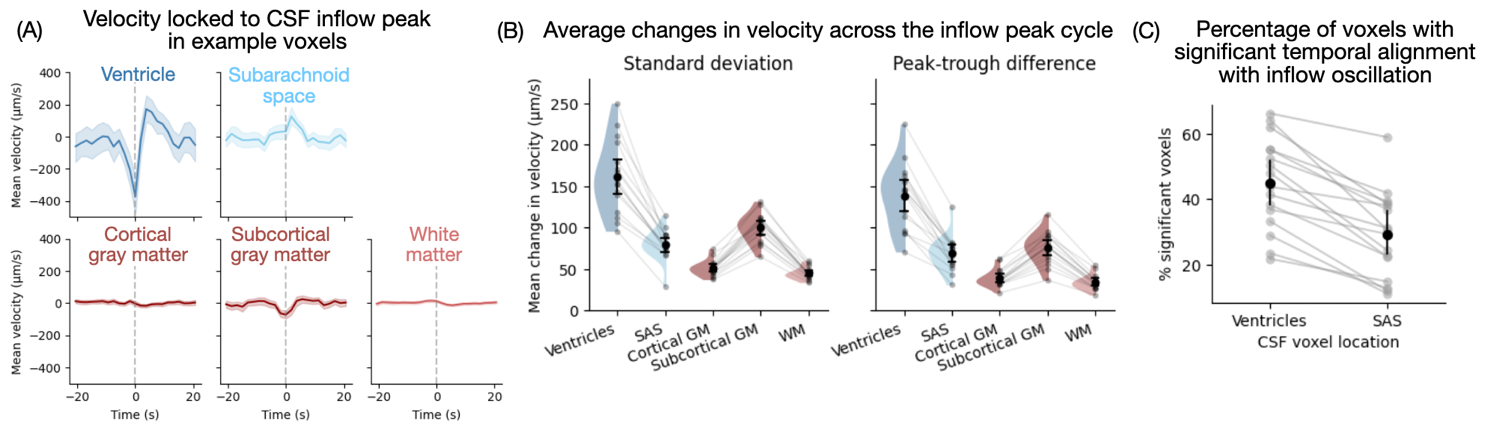

**Fig. S4.** **CSF flow velocity modulated by fourth-ventricle inflow.** (A) Example voxels showing CSF flow and tissue movement time-locked to the inflow peak. Shading indicates 95% CI. (B) Average changes in velocity across inflow peak cycle. The standard deviation and peak-to-trough difference were calculated for each voxel and averaged within ventricle, subarachnoid space, cortical gray matter, subcortical gray matter, and white matter masks. All pairwise differences between ROIs were significant (paired *t*-tests, *p* < 0.05). Gray lines and dots indicate subject averages. (C) Percentage of voxels showing significant temporal coupling with inflow peaks within subarachnoid space and ventricle masks. Dots indicate subject averages. Group-level means in black; error bars indicate 95% CI. SAS: subarachnoid space; GM: gray matter; WM: white matter.

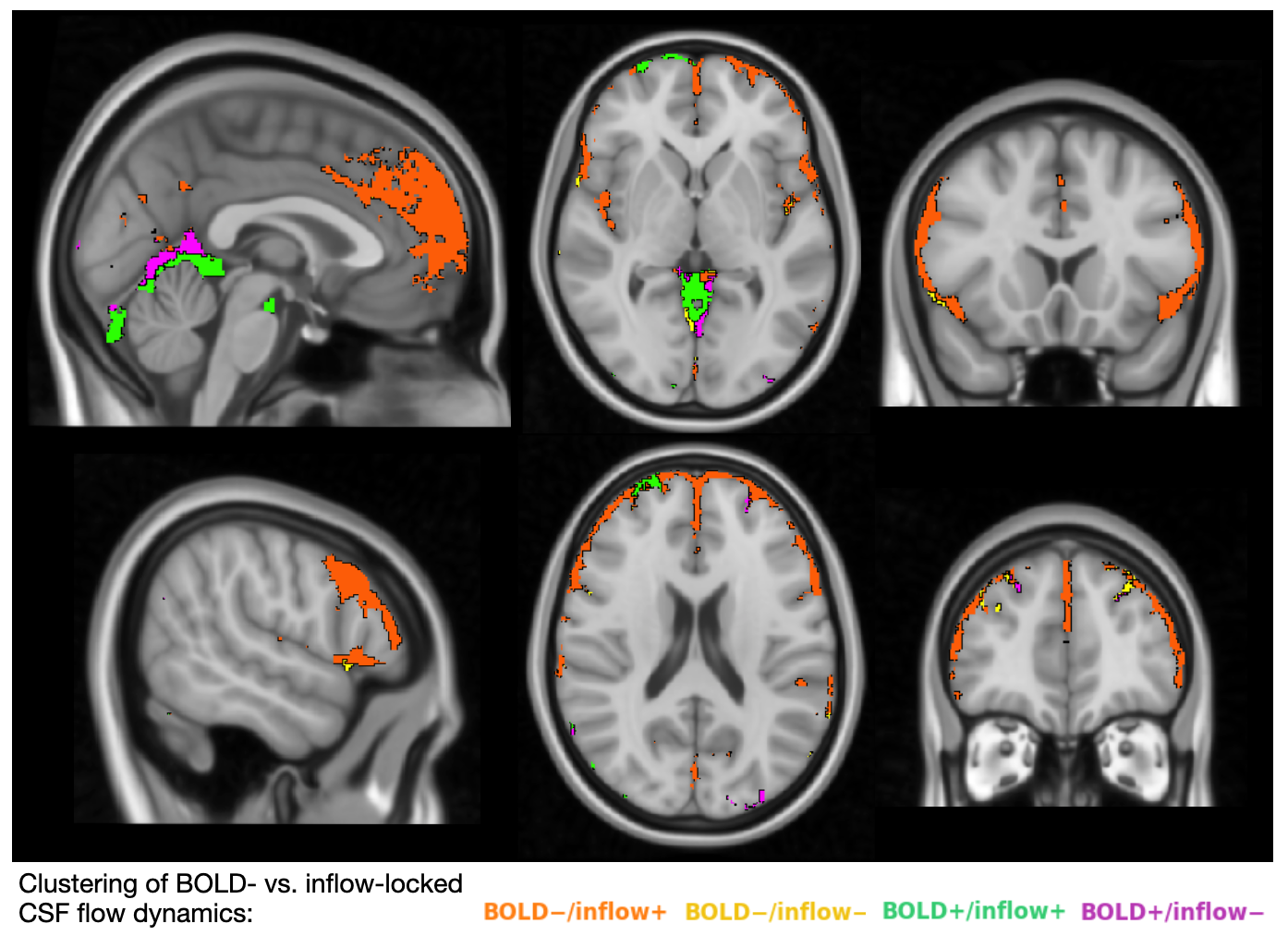

**Fig. S5. Spatial distribution of BOLD-locked vs. inflow-locked CSF coupling groups across the subarachnoid space.** The largest group, with negative BOLD-locked and positive inflow-locked coupling (BOLD−/inflow+; orange), covers most of the cortical-adjacent subarachnoid space and consists of CSF voxels whose flow reverses between the two events. This pattern is consistent with oscillatory motion driven by cortical hemodynamic activity, in which CSF moves one way around BOLD peaks and the other way around inflow peaks. The reverse combination (BOLD+/inflow−; purple) is more dispersed, likely reflecting local anatomical or geometric features that redirect flow relative to the dominant pattern. The two same-sign groups, which do not reverse between BOLD and inflow events, lie away from the cortical surface. Voxels with positive coupling to both events (BOLD+/inflow+; green) concentrate in the basal cisterns and move against net CSF drainage throughout the cycle, potentially facilitating circulation and mixing. Those with negative coupling to both (BOLD−/inflow−; yellow) appear in parts of the cisterns and in some regions near the cortex. Their patchier distribution makes them harder to attribute to a single pathway, and they may reflect a local driving force, such as proximity to large arteries or draining veins, that keeps flow moving one way.

**Comparison of Global and Local BOLD amplitude to predict CSF dynamics.**

To more directly dissociate global versus spatially specific contributions of BOLD amplitude to CSF dynamics, we compared linear mixed-effects models in which CSF outcomes were predicted by a global cortical BOLD amplitude measure alone, by a combined model including both global and local BOLD amplitude, and by a model additionally allowing the global × local interaction. For each outcome we fit linear mixed-effects models using maximum-likelihood estimation with scanning sessions as the grouping factor and ROI modeled as a variance component.

For CSF amplitude, the global-only model showed a robust positive association between global BOLD amplitude and CSF amplitude (β = 3.895, SE = 0.517, *z* = 7.536, *p* < 0.001). Adding local BOLD amplitude significantly improved model fit (likelihood-ratio test χ²_(1)_ = 56.482, *p* < 0.001), indicating that local BOLD amplitude explains variance in CSF amplitude beyond what is captured by the global measure. In this combined model, local BOLD amplitude remained strongly predictive of CSF amplitude (β = 1.847, SE = 0.246, *z* = 7.525, *p* < 0.001), whereas the global effect was no longer significant after accounting for local BOLD amplitude (β = −0.192, SE = 0.749, *z* = −0.256, *p* = 0.798). Allowing an interaction between global and local BOLD amplitude further improved model fit relative to the additive model (χ²_(1)_ = 16.038, *p* < 0.001). In the interaction model, the global × local term was negative (β = −1.951, SE = 0.487, *z* = −4.005, *p* < 0.001), suggesting that the positive association between local BOLD amplitude and CSF amplitude is attenuated as global BOLD amplitude increases.

For BOLD–CSF coupling strength, the global-only model showed a significant positive association between global BOLD amplitude and coupling strength (β = 0.017, SE = 0.003, *z* = 5.705, *p* < 0.001). Adding local BOLD amplitude significantly improved model fit (χ²_(1)_ = 14.779*, p* < 0.001). In this combined model, local BOLD amplitude was positive and significant (β = 0.006, SE = 0.001, *z* = 3.846, *p* < 0.001), whereas the global term was no longer significant (β = 0.005, SE = 0.004, *z* = 1.130, *p* = 0.258). The interaction model did not improve fit relative to the additive model (χ²_(1)_ = 0.545, *p* > 0.05).

Overall, these model comparisons support the conclusion that CSF amplitude is influenced by both global and local BOLD fluctuations, whereas BOLD–CSF coupling strength is more closely associated with local BOLD variation.

**Comparison of signal-to-noise ratio across bins**

Because local BOLD peak-locked events were sorted into bins based on the amplitude of the local BOLD oscillation, we tested whether the observed results could be explained by covariance between BOLD and CSF arising from noise rather than signal. To assess this, we quantified the mean-to-variability ratio of the event-locked CSF time series (CSF tSNR) and the corresponding cortical BOLD time series (BOLD tSNR) for each subject and bin. We then tested for differences across bins using a repeated-measures ANOVA with bin as a within-subject factor. If noise were the primary driver of the observed BOLD–CSF coupling, we would expect higher-amplitude bins (which showed stronger BOLD–CSF coupling strength) to also exhibit increased tSNR.

Both CSF and BOLD tSNR measures varied significantly across bins (CSF tSNR: *F*_(4,64)_ = 10.71, *p* < 0.001; BOLD tSNR: *F*_(4,64)_ = 17.13, *p* < 0.001). However, these differences were in the opposite direction of the noise-driven prediction: although local BOLD amplitude increased across bins, both CSF and BOLD tSNR decreased on average with increasing bin number. This could be driven by how the bins were characterized, namely by the maximum-minimum amplitude, which could itself be interpreted as a noise-related metric. This pattern argues against the interpretation that the positive association between BOLD amplitude and CSF dynamics is driven simply by higher signal-to-noise in higher-amplitude bins.

To further assess whether tSNR differences varied with arousal state, we examined CSF and BOLD tSNR separately for sleep and drowsiness across bins. Using repeated-measures ANOVA with both bin and state as within-subject factors, we found a significant main effect of state on CSF tSNR (*F*_(1,7)_ = 20.07, *p* < 0.01), but no significant effect of state for BOLD tSNR (*F*_(1,7)_ = 0.42, *p* = 0.54), suggesting that CSF signal is noisier during drowsiness than sleep. As a result, amplitude-based CSF measures are difficult to interpret across arousal states, because peak-to-trough estimates scale not only with the underlying signal but also with state-dependent differences in variance.

**Table S1. Linear mixed-effects models comparing BOLD-locked and cardiac-locked CSF dynamics.** Models were fit on ROI-level values from subarachnoid ROIs significantly coupled to each event type, with event type as a fixed effect and a random intercept for ROI.

**Velocity amplitude:**

| **Fixed effect** | **β** | **SE** | ***z*** | ***p*** | **95% CI** |
| --- | --- | --- | --- | --- | --- |
| Intercept | 174.547 | 13.043 | 13.383 | < 0.001 | [148.984, 200.110] |
| BOLD-locked vs. cardiac-locked | −155.210 | 18.056 | −8.596 | < 0.001 | [−190.599, −119.820] |

**Accumulated absolute displacement:**

| **Fixed effect** | **β** | **SE** | ***z*** | ***p*** | **95% CI** |
| --- | --- | --- | --- | --- | --- |
| Intercept | 47.071 | 8.818 | 5.338 | < 0.001 | [29.788, 64.354] |
| BOLD-locked vs. cardiac-locked | 116.049 | 11.188 | 10.373 | < 0.001 | [94.121, 137.976] |

**Directional consistency:**

| Fixed effect | β | SE | *z* | *p* | 95% CI |
| --- | --- | --- | --- | --- | --- |
| Intercept | 0.483 | 0.021 | 23.431 | < 0.001 | [0.443, 0.524] |
| BOLD-locked vs. cardiac-locked | −0.428 | 0.029 | −14.743 | < 0.001 | [−0.485, −0.371] |

**Table S2. Linear mixed-effects models for local BOLD amplitude predicting neighboring CSF amplitude and BOLD–CSF coupling.**

**CSF velocity amplitude:**

| **Dependent variable** | **Fixed effect** | **β** | **SE** | ***z*** | ***p*** | **95% CI** |
| --- | --- | --- | --- | --- | --- | --- |
| CSF amplitude | Intercept | 35.153 | 1.069 | 32.144 | < 0.001 | [32.995, 37.311] |
|  | Local BOLD amplitude | 1.802 | 0.169 | 10.663 | < 0.001 | [1.471, 2.133] |
|  | ROI | 1823.844 | 4.028 |  |  |  |

**BOLD–CSF coupling strength:**

| **Dependent variable** | **Fixed effect** | **β** | **SE** | ***z*** | ***p*** | **95% CI** |
| --- | --- | --- | --- | --- | --- | --- |
| Maximum absolute cross-correlation | Intercept | 0.650 | 0.009 | 72.470 | < 0.001 | [0.632, 0.668] |
|  | Local BOLD amplitude | 0.007 | 0.001 | 6.789 | < 0.001 | [0.005, 0.009] |
|  | ROI | 0.144 |  |  |  |  |

**Table S3. Linear mixed-effects models comparing global and local BOLD amplitude as predictors of CSF amplitude and BOLD–CSF coupling strength.** Global BOLD amplitude was defined as the mean cortical BOLD oscillation across all cortical regions, and local BOLD amplitude as the ROI-specific cortical BOLD oscillation for each matched tissue–CSF pair. Separate linear mixed-effects models were fit for CSF amplitude and BOLD–CSF coupling strength, including global BOLD alone, global plus local BOLD, and their interaction. Reported values are fixed-effect estimates (β), standard errors (SE), *z* statistics, *p* values, and 95% confidence intervals. Results indicate that CSF amplitude is influenced by both global and local BOLD fluctuations, whereas BOLD–CSF coupling strength is more closely associated with local BOLD variation. Model-comparison statistics reflect likelihood-ratio tests between nested models.

**CSF amplitude prediction model results:**

| **Model** | **Term** | **β** | **SE** | ***z*** | ***p*** | **95% CI** |
| --- | --- | --- | --- | --- | --- | --- |
| Global only | Intercept | 28.185 | 1.973 | 14.284 | < 0.001 | [24.318, 32.052] |
|  | Global BOLD amplitude | 3.895 | 0.517 | 7.536 | < 0.001 | [2.882, 4.908] |
| Global + Local | Intercept | 35.642 | 2.203 | 16.175 | < 0.001 | [31.323, 39.960] |
|  | Global BOLD amplitude | −0.192 | 0.749 | −0.256 | 0.798 | [−1.659, 1.276] |
|  | Local BOLD amplitude | 1.847 | 0.246 | 7.525 | < 0.001 | [1.366, 2.329] |
| Global × Local | Intercept | 12.433 | 6.198 | 2.006 | 0.045 | [0.285, 24.582] |
|  | Global BOLD amplitude | 6.697 | 1.875 | 3.571 | < 0.001 | [3.021, 10.372] |
|  | Local BOLD amplitude | 8.542 | 1.689 | 5.056 | < 0.001 | [5.231, 11.852] |
|  | Global × Local | −1.951 | 0.487 | −4.005 | < 0.001 | [−2.906, −0.996] |

**CSF amplitude prediction model comparison:**

| **Comparison** | **χ²** | **df** | ***p*** |
| --- | --- | --- | --- |
| Global + Local vs. Global only | 56.482 | 1 | < 0.001 |
| Global × Local vs. Global + Local | 16.038 | 1 | < 0.001 |

**BOLD–CSF coupling strength prediction model results:**

| **Model** | **Term** | **β** | **SE** | ***z*** | ***p*** | **95% CI** |
| --- | --- | --- | --- | --- | --- | --- |
| Global only | Intercept | 0.615 | 0.013 | 46.512 | < 0.001 | [0.589, 0.640] |
|  | Global BOLD amplitude | 0.017 | 0.003 | 5.705 | < 0.001 | [0.011, 0.023] |
| Global + Local | Intercept | 0.637 | 0.014 | 44.053 | < 0.001 | [0.609, 0.666] |
|  | Global BOLD amplitude | 0.005 | 0.004 | 1.130 | 0.258 | [−0.004, 0.014] |
|  | Local BOLD amplitude | 0.006 | 0.001 | 3.846 | < 0.001 | [0.003, 0.008] |
| Global × Local | Intercept | 0.612 | 0.037 | 16.365 | < 0.001 | [0.539, 0.685] |
|  | Global BOLD amplitude | 0.013 | 0.011 | 1.128 | 0.259 | [−0.009, 0.034] |
|  | Local BOLD amplitude | 0.013 | 0.010 | 1.289 | 0.197 | [−0.007, 0.033] |
|  | Global × Local | −0.002 | 0.003 | −0.738 | 0.460 | [−0.008, 0.004] |

**BOLD–CSF coupling strength prediction model comparison:**

| **Comparison** | **χ²** | **df** | ***p*** |
| --- | --- | --- | --- |
| Global + Local vs. Global only | 14.779 | 1 | < 0.001 |
| Global × Local vs. Global + Local | 0.545 | 1 | > 0.05 |

**Table S4. Linear mixed-effects models comparing BOLD–CSF coupling strength during sleep and drowsiness.** Top: voxel-wise coupling strength (maximum absolute cross-correlation between BOLD and CSF flow), using all available peak events, modeled separately for the ventricle and subarachnoid space (SAS compartments), with arousal state as a fixed effect and scan session as a random intercept. Bottom: a trial-matched replication of this comparison performed at the subarachnoid ROI level (see Fig. 3A), in which coupling strength was computed within each anatomically defined subarachnoid ROI and pooled across all ROIs; models were fit separately using the full sample of peak events and a trial-matched sample in which the arousal state with more available peak events was subsampled within each run (see Methods).

BOLD–CSF coupling strength

| **ROI** | **Fixed effect** | **β** | **SE** | ***z*** | ***p*** | **95% CI** |
| --- | --- | --- | --- | --- | --- | --- |
| Ventricle | Intercept | 0.481 | 0.014 | 34.182 | < 0.001 | [0.454, 0.509] |
|  | Drowsy vs. sleep | −0.056 | 0.018 | −3.043 | < 0.01 | [−0.091, −0.020] |
| SAS | Intercept | 0.493 | 0.019 | 26.152 | < 0.001 | [0.456, 0.530] |
|  | Drowsy vs. sleep | −0.067 | 0.019 | −3.529 | < 0.001 | [−0.105, −0.030] |

BOLD–CSF coupling strength, ROI-level analysis (full sample vs. trial-matched sample, across all ROIs)

| **Sample** | **Fixed effect** | **β** | **SE** | ***z*** | ***p*** | **95% CI** |
| --- | --- | --- | --- | --- | --- | --- |
| Full sample | Intercept | 0.682 | 0.018 | 38.507 | < 0.001 | [0.647, 0.717] |
|  | Drowsy vs. sleep | −0.063 | 0.019 | −3.271 | < 0.01 | [−0.100, −0.025] |
| Matched sample | Intercept | 0.686 | 0.019 | 35.980 | < 0.001 | [0.648, 0.723] |
|  | Drowsy vs. sleep | −0.051 | 0.016 | −3.142 | < 0.01 | [−0.082, −0.019] |

**Table S5. Linear mixed-effects models for local BOLD amplitude predicting neighboring CSF amplitude and BOLD–CSF coupling during sleep.**

**CSF velocity amplitude during sleep:**

| **Dependent variable** | **Fixed effect** | **β** | **SE** | ***z*** | ***p*** | **95% CI** |
| --- | --- | --- | --- | --- | --- | --- |
| CSF amplitude | Intercept | 41.738 | 1.371 | 30.439 | < 0.001 | [39.050, 44.425] |
|  | Local BOLD amplitude | 1.442 | 0.208 | 6.932 | < 0.001 | [1.035, 1.850] |
|  | ROI | 2837.197 | 5.260 |  |  |  |

**BOLD–CSF coupling strength during sleep:**

| **Dependent variable** | **Fixed effect** | **β** | **SE** | ***z*** | ***p*** | **95% CI** |
| --- | --- | --- | --- | --- | --- | --- |
| BOLD–CSF coupling | Intercept | 0.640 | 0.009 | 69.109 | < 0.001 | [0.622, 0.658] |
|  | Local BOLD amplitude | 0.008 | 0.001 | 7.567 | < 0.001 | [0.006, 0.010] |
|  | ROI | 0.152 |  |  |  |  |

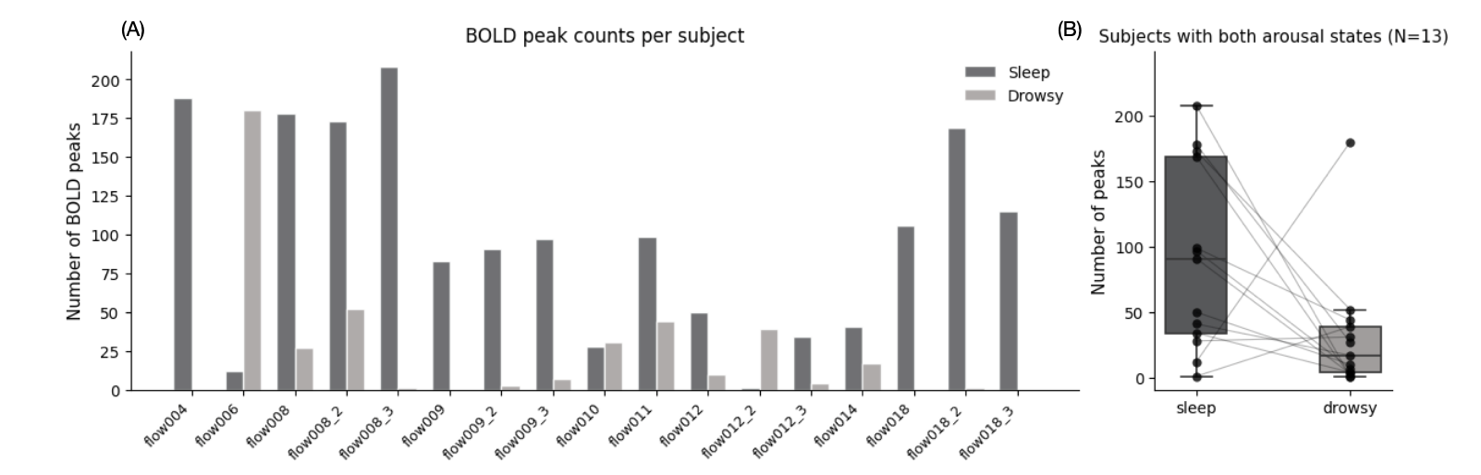
**Fig. S6. BOLD peak counts during sleep and drowsiness.** (A) Number of identified cortical BOLD peaks per subject, separated by arousal state (*N* = 17 sessions). Periods with ≤ 8 button presses per 20 s were categorized as sleep and periods with > 8 button presses per 20 s as drowsy. Across all subjects, 1673 peaks were identified during sleep and 416 during drowsiness. (B) Distribution of peak counts in scanning sessions with peaks in both arousal states (*N* = 13 sessions). Box plots show the median, interquartile range, and whiskers extending to the most extreme value within 1.5 × IQR; individual subjects are overlaid as connected dots. Subjects tended to have more peaks during sleep than drowsiness (paired *t*_(12)_ = 2.17, *p* = 0.051). Note that because scans began around midnight, elevated button press rates during behaviorally defined drowsy periods may reflect drowsy states rather than fully alert daytime wakefulness. Moreover, the substantially smaller number of detected peaks during drowsiness reduces the per-subject reliability of CSF velocity amplitude estimates. Because amplitude is computed as a peak-to-trough difference, it is sensitive to extreme values in the averaged waveform and is therefore inflated when fewer events are averaged per subject. This is consistent with the elevated CSF tSNR variability during drowsiness reported in *Comparison of signal-to-noise ratio across bins*. In contrast, BOLD–CSF coupling strength (absolute maximum cross-correlation) is a bounded, shape-based measure that is less sensitive to such variance differences. We therefore restrict main-text state comparisons to coupling strength rather than velocity amplitude (see Results).

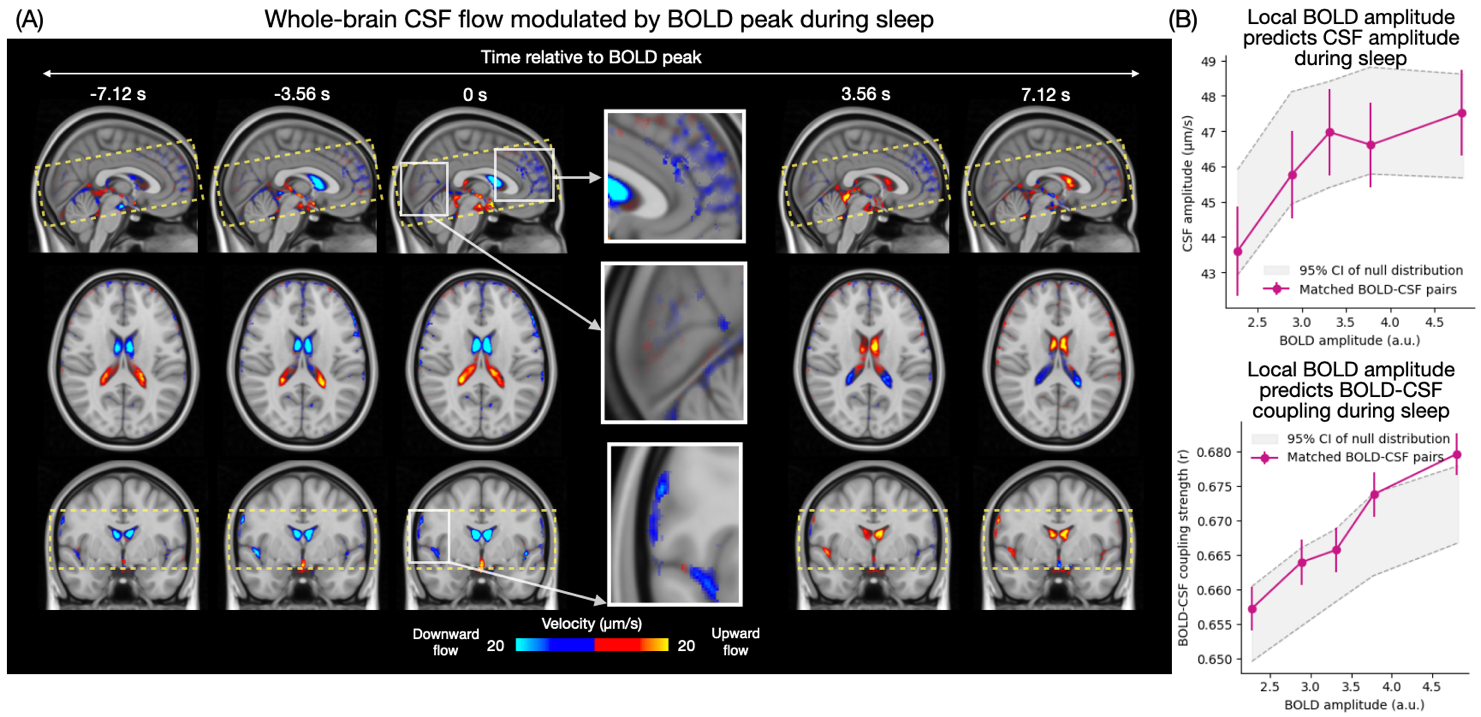

**Fig. S7. Replication of BOLD-locked CSF flow dynamics during sleep-only periods.** (A) Whole-brain CSF flow across BOLD peaks during sleep. CSF flow dynamics time-locked to BOLD peaks identified within sleep periods only were normalized and averaged across subjects. Yellow-to-red colors indicate upward flow, and blue voxels indicate downward flow. The whole-brain spatial pattern of CSF flow time-locked to BOLD peaks identified within sleep periods was visually consistent with the result using all periods (cf. Fig. 1E). To quantify this similarity, we computed both spatial and temporal correlations between voxel-wise BOLD-locked maps from the all-data and sleep-only analyses. The two sets of maps were highly correlated across space (mean Fisher-*z* transformed *r* = 0.98; one-sample *t*_(16)_ = 6.62, *p* < 0.001) and across time (mean Fisher-*z* transformed *r* = 0.98; one-sample *t*_(16)_ = 6.80, *p* < 0.001). (B) Quantification of regional BOLD–CSF relationships across matched ROI pairs during sleep. Larger local cortical BOLD amplitudes were associated with greater neighboring CSF velocity amplitude (top) and stronger local BOLD–CSF coupling strength, defined as the absolute maximum cross-correlation (bottom). Magenta lines indicate matched BOLD–CSF pairs, and the gray shaded region denotes the 95% confidence interval of the null distribution generated by randomly permuting tissue–CSF pairings. Repeating the binned linear mixed-effects analysis using sleep-only data, we found that local BOLD amplitude remained positively associated with both neighboring CSF velocity amplitude (β = 1.44, *p* < 0.001; permutation-based two-sided *p* < 0.01) and BOLD–CSF coupling strength (β = 0.008, *p* < 0.001; permutation-based two-sided *p* < 0.001; Table S5). These results indicate that the spatial pattern of BOLD-locked CSF dynamics and the regional dependence of CSF dynamics on local BOLD fluctuations are preserved within sleep periods alone.

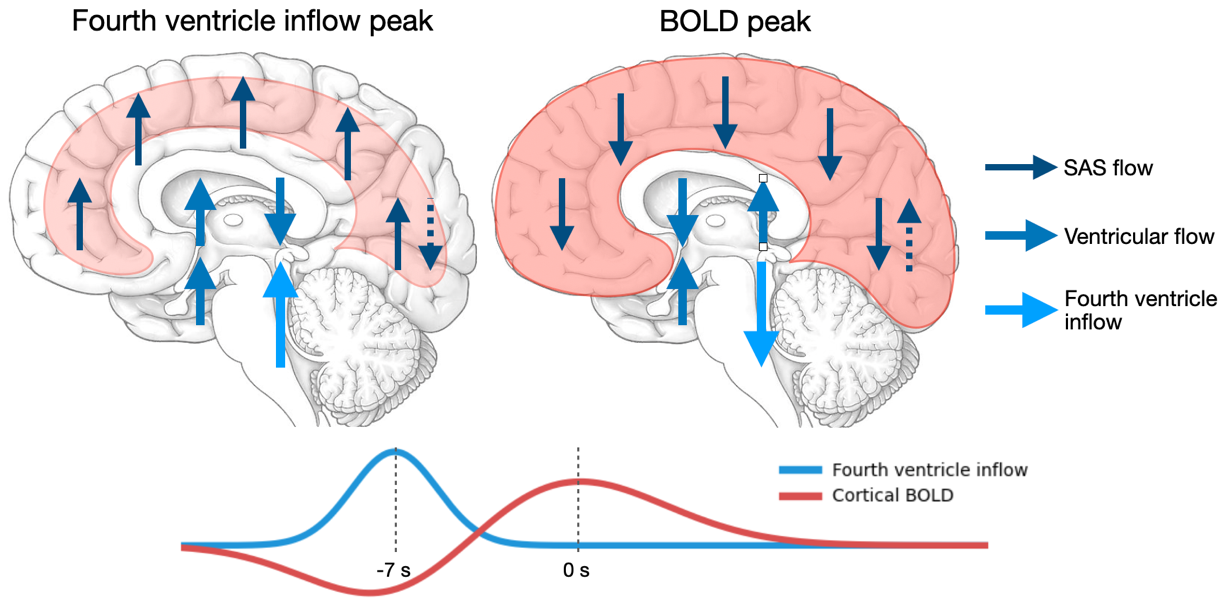

**Fig. S8. Schematic of the proposed brain-wide CSF cycle during sleep.** Fourth-ventricle inflow peaks and cortical BOLD peaks reflect two phases of the same continuous oscillation. Sagittal schematics show CSF flow direction at each phase. At the inflow peak, cortical blood volume decreases and CSF enters the fourth ventricle; subarachnoid CSF flows predominantly upward toward the dorsal convexity; the lateral ventricles show downward flow in the posterior segment and upward flow in the anterior segment; and CSF in most basal cistern regions flows upward. At the BOLD peak, approximately 7 s later, cortical blood volume increases and the pattern reverses: subarachnoid CSF flows predominantly downward, the lateral ventricles show upward flow in the posterior segment and downward flow in the anterior segment, and CSF in the fourth ventricle flows downward, whereas CSF in most basal cistern regions continues to flow upward. Illustration at the bottom shows the fourth-ventricle inflow and cortical BOLD time courses. Arrows indicate flow fluctuations relative to mean flow and show the dominant pattern across regions; solid arrows indicate the dominant direction, and dashed arrows indicate the less-common, opposite-direction flow in the subarachnoid space. Arrow color indicates compartment (dark blue: subarachnoid; medium blue: ventricular; light blue: fourth-ventricle inflow). Red shading indicates changes in cortical blood volume.
